# Functional architecture of intracellular oscillations in hippocampal dendrites

**DOI:** 10.1101/2024.02.12.579750

**Authors:** Zhenrui Liao, Kevin C. Gonzalez, Deborah M. Li, Catalina M. Yang, Donald Holder, Natalie E. McClain, Guofeng Zhang, Stephen W. Evans, Mariya Chavarha, Jane Yi, Christopher D. Makinson, Michael Z. Lin, Attila Losonczy, Adrian Negrean

## Abstract

Fast electrical signaling in dendrites is central to neural computations that support adaptive behaviors. Conventional techniques lack temporal and spatial resolution and the ability to track underlying membrane potential dynamics present across the complex three-dimensional dendritic arbor *in vivo*. Here, we perform fast two-photon imaging of dendritic and somatic membrane potential dynamics in single pyramidal cells in the CA1 region of the mouse hippocampus during awake behavior. We study the dynamics of subthreshold membrane potential and suprathreshold dendritic events throughout the dendritic arbor *in vivo* by combining voltage imaging with simultaneous local field potential recording, *post hoc* morphological reconstruction, and a spatial navigation task. We systematically quantify the modulation of local event rates by locomotion in distinct dendritic regions and report an advancing gradient of dendritic theta phase along the basal-tuft axis, then describe a predominant hyperpolarization of the dendritic arbor during sharp-wave ripples. Finally, we find spatial tuning of dendritic representations dynamically reorganizes following place field formation. Our data reveal how the organization of electrical signaling in dendrites maps onto the anatomy of the dendritic tree across behavior, oscillatory network, and functional cell states.

## Introduction

Behavior-contingent network oscillations coordinate active neural ensembles in cortical circuits^1–6^. Cortical neurons primarily interface with behavioral state-dependent population patterns at their dendritic arbor, receiving thousands of afferent inputs along hundreds of microns^7^. Distinct morpho-electrical properties of these small-diameter, highly branched dendrites support dynamic compart-mentalization for synaptic integration and plasticity, spike initiation and propagation, and intrinsic oscillatory dynamics^8–12^. How these dendritic processes interact to produce behaviorally-relevant single-cell computations *in vivo* remains a fundamental open question in the field of neuroscience. Due to technical limitations, it has previously not been possible to interrogate dendritic membrane potential dynamics in different behavior states and network oscillations. Cortical dendrites are seldom accessible to direct electrophysiological recordings *in vivo*^13–15^, and have primarily been studied either *in vitro* or with optical imaging using calcium sensors *in vivo*^14;16–19;19–24^. Direct electrophysiological recordings are particularly challenging from small-diameter dendrites, even *in vitro*^25;26^. While informative, calcium imaging lacks the temporal resolution and fidelity to membrane potential dynamics required to answer this question.

Genetically encoded voltage indicators (GEVIs) have recently emerged as promising tools for fast optical recording of neural circuit dynamics^27–30^. To study the relationship between subcellular voltage dynamics and network states *in vivo*, we electroporated ASAP3^28^ into single pyramidal cells and performed two-photon voltage imaging across the somato-dendritic membrane surface of pyramidal cells in the CA1 area of the hippocampus (CA1PCs), where dendritic processing is critical for spatial navigation and episodic memory^5;8;17;31–35^. Using concurrent hippocampal local field potential (LFP) recordings, we relate dendritic voltage dynamics to two physiologically antagonistic population patterns: the locomotion-associated theta rhythm (5–10 Hz) and the immobility-associated high-frequency sharp-wave ripple (SWR) oscillation, which respectively typify memory encoding and consolidation-related network states in the hippocampus^6;36;37^. Our results reveal dynamic compartmentalization of intracellular voltage signals depending on behavioral, network, and cellular states within the dendritic arbor of CA1PCs *in vivo*.

## Results

### Two-photon voltage imaging dendritic activity *in vivo*

To investigate electrical signaling in dendrites of CA1PCs in behaving mice, we performed *in vivo* single-cell electroporation (SCE) under two-photon microscopy guidance^38–40^ to deliver plasmids for stable expression of ASAP3 (Fig S1), a green-fluorescent GEVI developed and characterized for use *in vivo*^28^, along with mRuby3^41^, a static red-fluorescent morphological filler protein (see Methods). Simultaneous two-photon ASAP3-voltage imaging of CA1PCs and contralateral LFP recordings^42^ were performed in head-fixed mice during volitional running and immobility on a cued treadmill^43^ (Figs 1a, S2a,b, see Methods). Our SCE procedure allowed for bright labeling of fine dendrites and *post hoc* full reconstruction of cell morphology (Fig 1b). In this work, we introduce phylogenetic dendrograms (“phylograms”) to render 3D dendritic morphology in two dimensions (Fig 1b, right; also see Methods). ASAP3 fluorescence was sampled at a frame rate of 440 Hz over a period of 30–45 s from 28–64 (median: 43, IQR: 10.75) regions of interest (ROIs) along the dendritic arbor (Figs 1c, S2c). Our labeling and recording configuration allowed for detection of depolarizing events (DEs, Fig 1c) using a matched-filter approach (Figs S2d,e, S3-S5).

**Figure 1.**
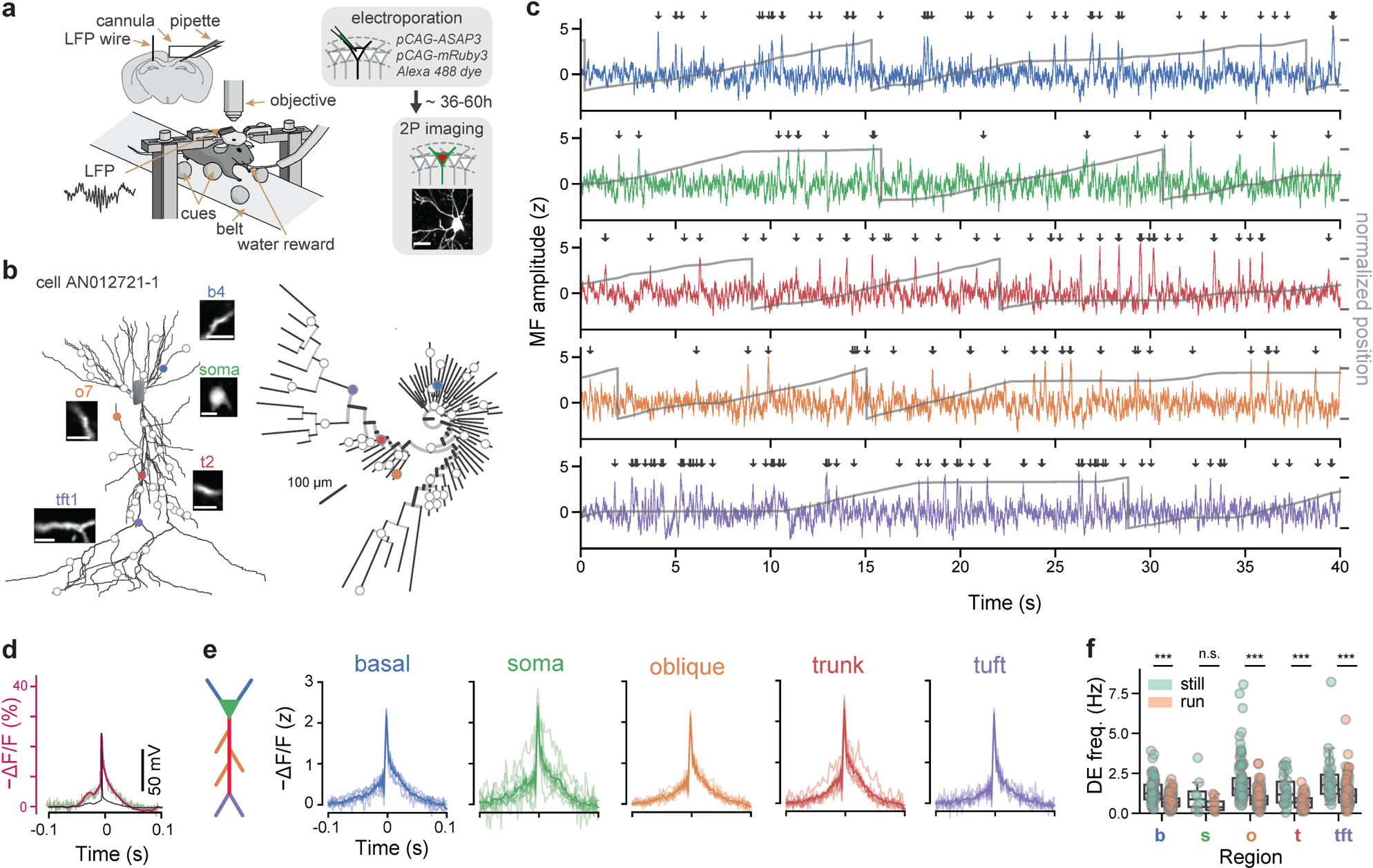
Detection and organization of depolarizing events (DEs) across the dendritic arbor and behavioral states. **a.** Experimental setup. Electroporation-imaging cannula-window was implanted above the right dorsal hippocampal area CA1. A tungsten wire LFP electrode was inserted contralaterally, within or in proximity to the CA1 cell body layer. Mice were habituated using water rewards to run on a cued belt treadmill. ASAP3 and mRuby3 plasmids were single-cell electroporated (SCE) in CA1 pyramidal cells under two-photon microscopy guidance. Two-photon fluorescence measurements were obtained after approx. 36 hours post SCE from several tens of dendritic regions of interest (ROIs) across the whole dendritic tree. Scale bar 25 *µ*m. **b.** Morphological reconstruction and identification of recording locations. Left: two-photon microscopy stacks were acquired *in vivo* following each imaging session and neuronal morphologies were reconstructed (example cell AN012721-1). Inset scans show the extent of dendritic (basal: *b4*, trunk: *t2*, oblique: *o7*, tuft: *tft1*) and somatic imaging regions (scale bars 10*µ*m). Right: phylogram representation of the same cell’s branching morphology, with radial distance from soma representing path distance to each recording location. Circles indicate imaging locations. **c.** Example optical measurements of sequentially recorded dendritic electrical activity from cell shown in **b.** during bouts of locomotion and rest. Optical signals were sampled at 440 Hz, stabilized against movement artifacts during locomotion epochs, match-filtered (MF) and thresholded for depolarizing event (DE) detection (arrows) (see Methods). Note that ASAP3 fluorescence is plotted as −ΔF/F to reflect the direction of change in the membrane potential. Grey traces: animal position during the recording. **d.** Electrophysiological calibration: A somatically recorded CA1PC action potential (black) was played through voltage-clamped ASAP3-expressing HEK293 cells at 37 °C and the corresponding ASAP3 fluorescence response (green) was recorded. Magenta: Markov model predicted ASAP3 response. **e.** Average z-scored DE waveforms for isolated events pooled from different dendritic domains with a fluorescence baseline SNR >10 (calibrated against a shot-noise baseline) during locomotion and immobility behavior states (N=13 cells from 9 mice, colored curves). **f.** DE frequency is reduced during locomotion states compared to immobility across all dendritic domains for fluorescence baseline SNR >10 (basal: *p* = 10^−10^, oblique: *p* = 2 × 10^−15^, soma: *p* = 0.03, trunk: *p* = 2 × 10^−6^, tuft: *p* = 4 × 10^−5^, Wilcoxon paired test; 587 dendrites from 13 cells). Two-way ANOVA for frequency: *p* = 3 × 10^−11^, main effect of region; *p* = 1 × 10^−41^, main effect of locomotion; *F* (4, 770) = 2.24*, p* = 0.06 region×locomotion interaction. Box: Q1, Q2, Q3; whiskers: range (excluding outliers). Point represent individual dendrites. *α* = 0.01 indicates significance for *post hoc* tests at the per-region level using Bonferroni’s correction.

To better understand the voltage-to-fluorescence transformation of ASAP3, we modeled sensor dynamics using a 4-state Markov chain based on recordings from ASAP3-expressing voltage-clamped HEK293 cells (Figs 1d, S3, see Methods). We then compared the model-predicted ASAP3 response to single action potentials recorded *in vivo* to detected DEs in our dataset, and found close qualitative agreement (Fig 1d, see Methods). The average DE waveform detected using our method was qualitatively similar across dendritic domains Fig 1e and between running and immobile behavior states (Figs S2b, S6). Across behavior states, the soma fired at an average of 0.67 ± 0.14 Hz. Mean DE frequency differed significantly between dendritic regions (basal: 0.89 ± 0.03 Hz, trunk: 0.93 ± 0.08, oblique: 1.1 ± 0.04, tuft: 1.4 ± 0.08; Kruskal-Wallis *H*-test, *H* = 37.8*, p* = 3 × 10^−8^) but did not vary as a function of distance along the basal-apical axis (Fig S6a). DE frequency decreased across all cellular domains during locomotion compared to rest (Fig 1f). These results demonstrate that DEs with fast sodium spike-like kinetics are detectable with voltage imaging across behavioral states and dendritic regions of CA1PCs *in vivo*. Throughout the remainder of this work, DEs refer to fast sodium spike-like events, likely representing both locally generated and backpropagating action potentials in dendrites, detected using a matched-filter based on a template calibrated to simultaneous patch-clamp and voltage imaging in an *in vitro* preparation (Methods). We also observed a smaller proportion of longer duration dendritic events in all dendritic regions (10.4% of all events, 56.2 ± 2.2 ms, mean ± s.e.m.), likely representing sodium spikes with a slower, NMDA receptor/voltage-gated calcium channel-mediated component^44;45^ using a template constructed from dendritic patch-clamp recordings from CA1 acute slices (Fig S7a)^46^. Our fast voltage imaging approach necessitates sequential scanning of ROIs, which does not allow for disambiguation of backpropagating action potentials from locally generated dendritic spikes or an exhaustive characterization of the diversity of dendritic events. In the remainder of this work, we characterize dendritic signalling in the online (locomotor) and offline (immobile) states using subthreshold membrane voltage and suprathreshold fast DEs.

### Theta oscillation-related modulation of dendritic membrane potential

When an animal is moving, a 5–10 Hz extracellular oscillation (“theta”) emerges in the hippocampus, which is thought to reflect the “online” state of the brain and support memory encoding and decoding^37^. Layer-dependent organization of theta has been proposed based on extracellular recordings and recordings in anesthetized animals^37^. Here, we investigated the relationship between extracellular theta and intracellular membrane potential across the entire dendritic tree in the awake, behaving animal. Previous work has found high coherence between interhemispheric hippocampal theta oscillations^47;48^; in order to avoid electrical artifacts induced by imaging, we imaged membrane potential while simultaneously recording LFP in contralateral CA1 (see Methods). Out of *N* = 11 cells, ASAP3 fluorescence from 83% basal, 100% somatic, 97% trunk, 91% oblique and 93% tuft ROIs was found to be theta-band modulated (example cell in Fig 2a, see Methods). Morphological reconstruction revealed that theta oscillation amplitude and phase changed gradually across the dendritic tree as a function of somatic path distance to the recorded ROI (example cell in Fig 2b, top, middle, Fig 2c, additional examples in Fig S8). Regression analysis of imaged ROIs within the same cell showed a significant gradient of theta-band fluorescence oscillation phase relative to the somatic compartment along the basal-tuft axis for 8/11 cells, with 7/8 showing an advancing gradient toward the tuft (Fig 2d, top, middle). The average theta-band fluorescence oscillation amplitude of theta-modulated ROIs was also found to increase from basal toward the tuft domain (Fig 2d, bottom). Pooling ROIs across cells, starting from the distalmost basal and progressing toward distalmost tuft dendritic ROIs, fluorescence oscillation phase advanced along the basal-tuft axis with a slope of −7.9^°^/100*µ*m while oscillation amplitude increased with a slope of = 0.7%*z*/100*µ*m (Fig 2e).

**Figure 2.**
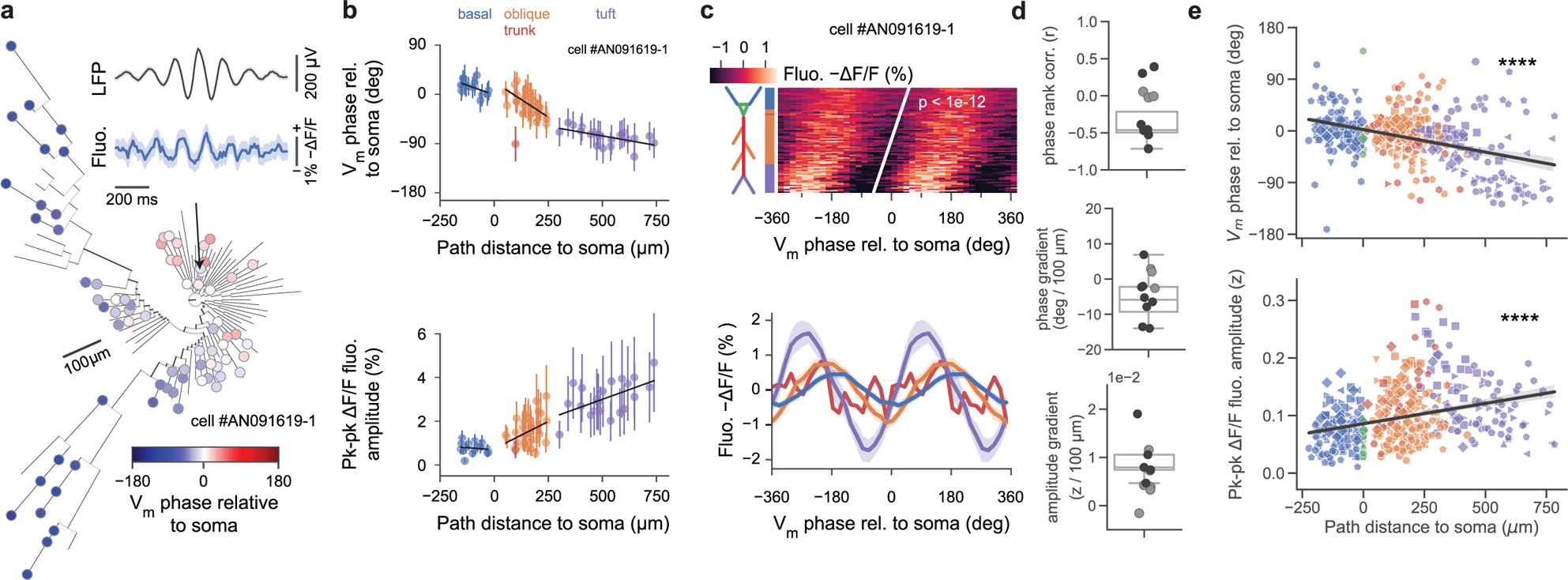
Theta oscillation is organized across the dendritic tree as a traveling wave. **a.** Phylogram of fluorescence phase at recording locations, relative to estimated somatic phase (all regions showed significant sinusoidal phase modulation at *p <* 0.05). Inset, top: Contralaterally-recorded extracellular theta-band-filtered (5–10 Hz) local-field potential; Bottom: Cycle-averaged fluorescence oscillation of a basal dendritic segment (arrow, mean with 95% CI). **b.** Dendritic tree gradients in theta oscillation phase, amplitude and DE phase for example cell in **a.** during locomotion epochs. Top: Membrane potential oscillation phase advances along the basal-tuft axis relative to somatic phase (zero phase corresponds to a trough in somatic membrane potential oscillation). Region fits: phase dependence: basal (*r*^2^ = 0.27*, p* = 0.04), oblique (*r*^2^ = 0.39*, p <* 0.01), tuft (*r*^2^ = 0.32*, p* = 0.01); overall (*r* = −0.89*, p <* 6 × 10^−27^). Bottom: Membrane potential oscillation amplitude increases along the basal-tuft axis. Region fits: basal (*r*^2^ = 0.02*, p* = 0.62), oblique (*r*^2^ = 0.13*, p* = 0.02), tuft (*r*^2^ = 0.30*, p* = 0.01); overall (*r* = 0.84*, p <* 2 × 10^−21^). **c.** Subthreshold theta-band membrane potential oscillation is a traveling wave across the basal-tuft dendritic tree axis. Top: Heatmap of mean phase-binned fluorescence vs soma path-distance rank along basal-tuft axis for example cell in **a.** (two cycles shown) with rank-regression line (*r* = −0.70*, p <* 10^−12^). Vertical axis colored by cell region. Bottom: Cycle-averaged mean fluorescence of example cell as a function of theta phase relative to somatic phase. **d.** Top: Correlation coefficients from rank-regression shown in **c.** (top) of path-distance to soma vs phase bin of minimum. 8/11 cells recorded exhibit a significant phase gradient along the basal-tuft axis (black dots: significant cells; grey dots: nonsignificant cells), with 6/8 having a negative (advancing) gradient. Middle: Phase gradients from least-squares fit as in **e.** calculated per-cell. 8/11 cells recorded show a significant phase gradient along the basal-tuft axis, with 7/8 having a negative (advancing) gradient. Bottom: Z-score amplitude gradients from least-squares fit as in **e.**calculated per-cell. 5/11 cells recorded exhibit a significant increasing gradient along the basal-tuft axis. **e.** Population summary of theta-band membrane potential oscillation phase and amplitude gradients by recording segment during locomotion epochs. (colors: regions, markers: *N* = 11 unique cells). Top: Theta-band membrane potential oscillation phase advances with distance from basal to tuft regions (slope = −7.9°/100*µ*m, Pearson’s r: *r* = −0.41*, p* = 1 × 10^−19^, Spearman’s rho: *ρ* = −0.35*, p* = 2 × 10^−14^). Bottom: Theta-band membrane potential oscillation z-score amplitude increases with distance from basal to tuft regions. (slope = 0.7%*z*/100*µ*m, Pearson’s r: *r* = 0.32*, p <* 8^−13^, Spearman’s rho: *ρ* = 0.37*, p <* 1 × 10^−16^)

### Theta phase preference of dendritic events

Somatic spikes occur preferentially at the extracellular theta trough, corresponding to the peak of the intracellular theta oscillation^37^. We asked if the same was true in dendrites. We found that gradients in theta phase and amplitude were accompanied by a comparable gradient in DE phase preference (example cell in Fig 3a-c, resultant DE phase distributions in Fig 3d), with DE phase advancing toward the tuft with a slope of −7.8^°^/100*µ*m (Fig 3e). A strong correlation between theta-band membrane-potential oscillation and DE phases was observed (Fig 3f), with the oscillation phase showing a slight advancement compared to DE phase, indicating that at the segment level, DEs occur preferentially close to the peak of the subthreshold oscillation. We expect locally-generated events to be entrained to local theta phase (45°slope of the DE phase vs membrane potential phase relationship) and bAPs to occur preferentially at the peak of somatic theta (0°slope). The true observed slope is shallower than 45°, consistent with a mixture of local events and somatic-phase conveying back-propagating action potentials. Unlike fast DEs, the distribution of slow events appear to be concentrated around 0°in all regions but the tuft (Fig S7b). To assess whether DE properties are modulated by the subthreshold oscillation, we performed a linear-circular regression of DE peak amplitude against DE phase (see Methods). DE amplitude was not strongly modulated by theta phase in any regions other than the tuft (*p <* 4 × 10^−5^, Fig S9).

**Figure 3.**
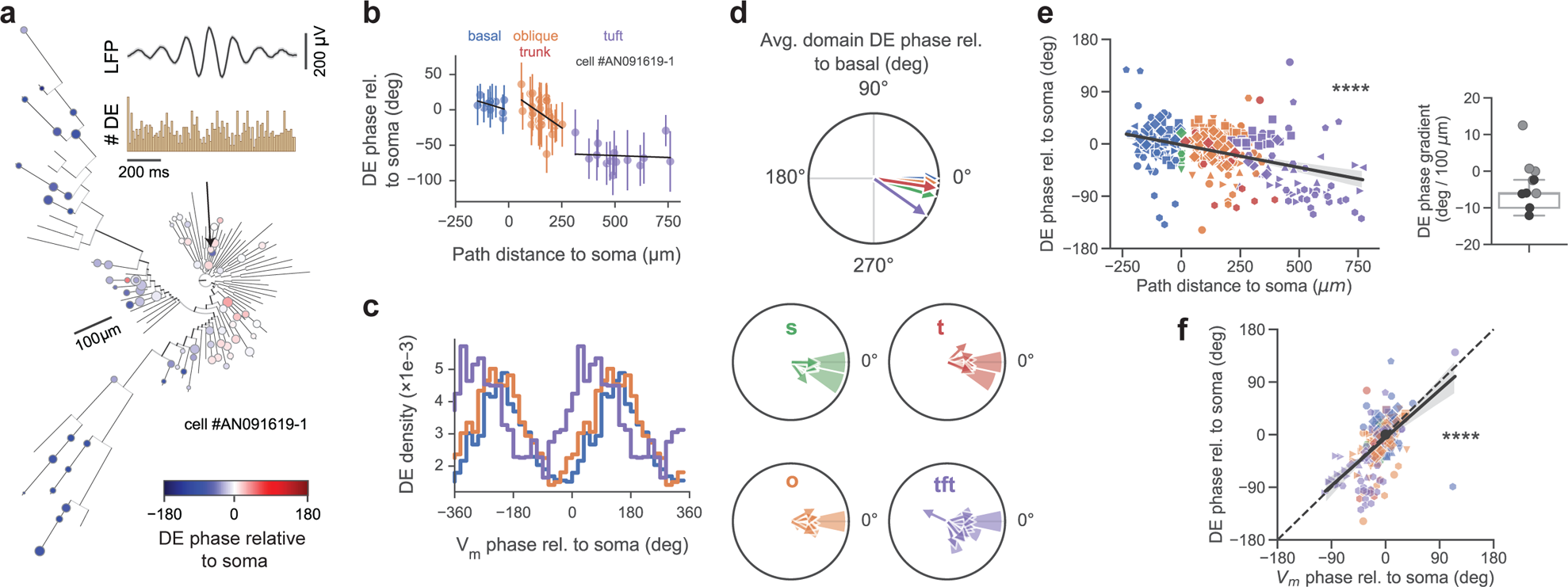
Depolarizing events are organized across the dendritic tree by local theta oscillations. **a.** Phylogram of depolarizing event (DE) phase at recording locations, relative to estimated somatic phase (all regions showed significant phase modulation at *p <* 0.05). Inset, top: Contralaterally-recorded extracellular theta-band-filtered (5–10 Hz) local-field potential; Bottom: DE histogram, with DEs occurring preferentially at the peak of intracellular theta oscillation. **b.** Dendritic tree gradient in DE phase for example cell in **a.** during locomotion epochs. DE preferred phase advances along the basal-tuft axis relative to somatic DE phase. Region fits: basal (*r*^2^ = 0.18*, p* = 0.2), oblique (*r*^2^ = 0.2*, p* = 0.02), tuft (*r*^2^ = 0.01*, p* = 0.79); overall (*r* = −0.81*, p <* 4 × 10^−13^). **c.** Histogram of DE densities for same example cell vs theta phase in the basal, oblique, trunk, and tuft domains relative to somatic membrane potential oscillation phase (two cycles shown, zero phase corresponds to estimated trough of somatic theta). DE density was defined as the DE rate (events / second) in each phase bin. **d.** Resultant phase of DEs (relative to basal mean phase) also changed gradually along the basal-tuft axis. ROI preferred phases (histograms) and cell means (arrows) by region. **e.** Left: Theta-band DE phase preference gradient across the basal-tuft axis relative to soma (slope = −7.8°/100*µ*m, *r* = −0.41*, p <* 10^−14^). Right: Strength of DE phase gradient by cell (significant cells in black, 5/10; boxplot significant only). **f.** Dendritic DEs preferentially occur close to the peak of local theta-band dendritic membrane potential oscillations. Comparison of DE resultant phase vs theta-band membrane potential oscillation phase, by recording segment (slope = 0.91 ± 0.07 (mean ± s.e.m.), (0.77, 1.04) 95% CI, *r* = 0.59*, p <* 10^−30^; colors: regions; markers: cells). Unless otherwise specified, all box-and-whisker plots show Q1, Q2, Q3, and range excluding outliers; points represent individual cells; *r* values are Pearson’s r; using Bonferroni correction *α* = 0.01 for *post hoc* tests performed on the per-region level, *α* = 0.05 for all other tests.

We then asked whether these gradients were explained by absolute distance from the soma or distance along the basal-apical axis. We focused on trunk and oblique segments, decomposing the total somatic path distance into an along-trunk component and an away from the trunk (oblique) component. Multivariate regression against these two components revealed that the observed gradients did not advance significantly along radial obliques dendrites; rather, they could be explained solely by the distance along the soma-trunk axis (Fig S10). Taken together, these results form the first subcellular characterization of theta oscillations across the CA1PCs dendritic tree in the behaving animal, and reveal strong correlation between the phase of local dendritic events and dendritic membrane theta phase.

### SWR-related modulation of dendritic membrane potential

Hippocampal dynamics during immobility (the “offline” state) differ considerably from online dynamics during locomotion. The theta oscillation disappears when an animal is immobile, while the sharp-wave ripple (SWR) emerges. SWRs are critical for offline memory consolidation, but like theta, have thus far primarily been studied at the somatic compartment. Here, we examine dendritic signalling across the arbor inside and outside of SWRs^36^. Overall, dendrites fired at 0.50 ± 0.08 Hz inside of SWRs compared to 1.50 ± 0.17 Hz (mean ± s.e.m.) during immobility outside of SWRs. On average, 17.6 ± 3% of each cell’s dendrites (mean ± s.e.m., *N* = 8 cells) were significantly modulated by SWRs; 98.3% of modulated dendrites showed a reduction in DE rate inside SWRs compared to immobility baseline outside of SWRs (example cell in Fig 4a, more cells in Extended Data Fig S11). In a ±500 ms peri-SWR window, we found a net suppression of dendritic DEs, which begins approximately 200 ms prior to SWR onset, reaching a nadir shortly after the peak of SWR power, and returning to baseline 200 ms after SWR peak (Fig 4b). We dissected this effect by dendritic region, and found that SWRs on average are associated with an acute hyperpolarization of dendritic membrane potential in every region, which reaches a nadir shortly after the SWR LFP power peak (Fig 4c, top). However, individual SWR responses are highly heterogeneous; while the predominant effect is inhibition, a minority of SWRs recruit the recorded dendrite (Fig S12). In each region, fluorescence showed maximal membrane potential hyperpolarization that was delayed from the SWR LFP peak power (baseline=0% − Δ*F/F*, Fig 4d, e, Tables S4, S5). The timing of this effect was fairly consistent between cells and along the basal-tuft axis (Fig S13). Concomitantly, we found an acute reduction in DE rate in the soma and all regions of the arbor (Fig 4c,bottom, Fig 4f, Table S6). Similar to fast DEs, slow events were also suppressed around SWRs (Fig S7c). Given that different subtypes of GABAergic interneurons are differentially recruited to SWRs and target distinct dendritic domains^49^, we asked whether DEs occurring during SWRs differed from those occurring outside SWRs in each region. Unlike somatic action potentials, local dendritic events are not stereotyped or all-or-none, but vary in their amplitude and morphology^44;50–55^ which may be modulated by oscillations^53;56^. We found that DE amplitudes were significantly decreased (Fig 4g) inside SWR epochs for the basal domain (outside: 18.9% −Δ*F/F*, 95% CI [18.1,19.5], inside: 15.7% −Δ*F/F*, 95% CI [13.9,18.3]; Fig 4h, Table S7), while amplitude was preserved but DE width was significantly narrowed in the oblique domain (outside: 26.0 ms, 95% CI [20.0, 30.1]; inside: 16.1 ms 95% CI [12.6, 21.7]; Fig 4i, Table S8); other regions did not exhibit significant differences in these parameters. In summary, we found that SWRs on average are associated with an inhibitory effect on the dendritic arbor, in membrane potential, DE rate, and DE waveform shape in the basal and oblique domains. This cell-wide inhibition begins preceding the peak of SWR power, peaks shortly after SWR LFP power peak, and recedes about 200 ms after the peak.

**Figure 4.**
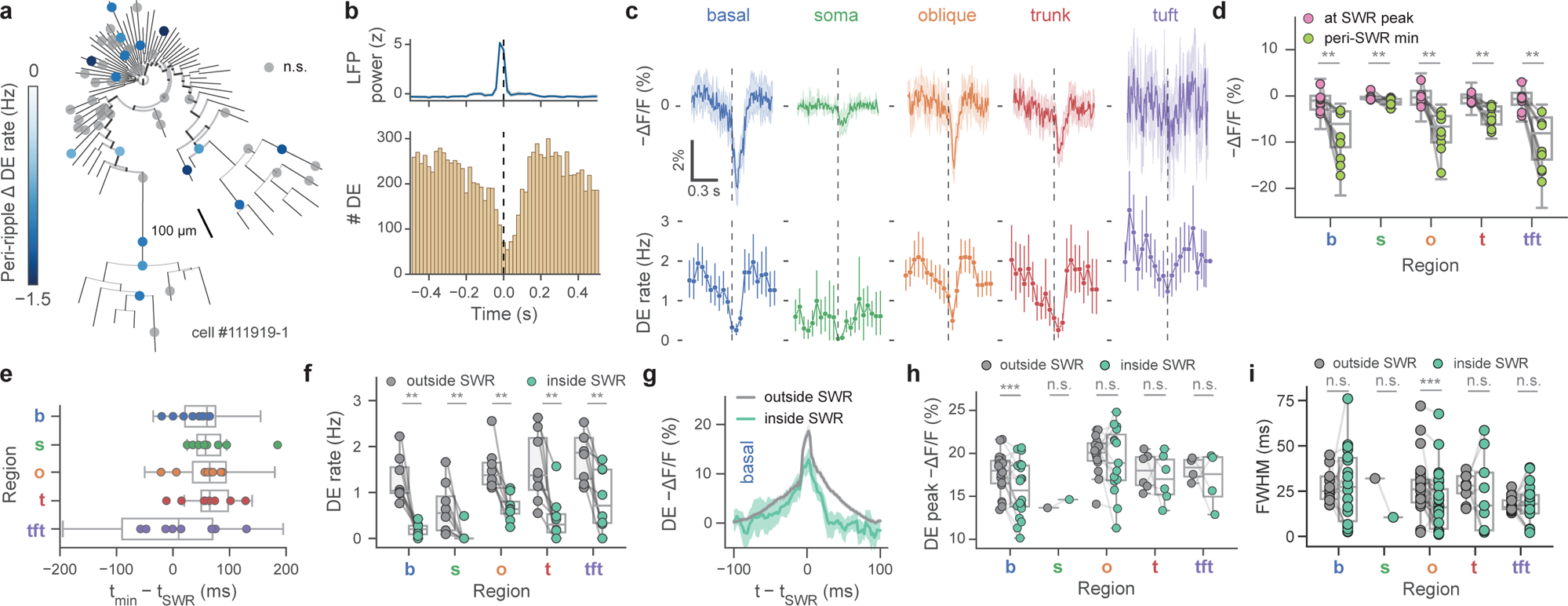
Sharp-wave ripple-associated modulation of dendritic electrical signaling. **a.** Example phylogram of a reconstructed cell (cell #111919-1) with significantly SWR-modulated recording locations colored by DE rate change inside minus outside of ripple epoch (grey: *p >* 0.05, two-tailed Poisson rate test). Significantly modulated recording locations show a suppression of DE rate during SWRs. **b.** Top: Mean z-scored LFP power in ripple band. Bottom: Peri-SWR spike time histogram (all regions, *N* = 8 cells). **c.** SWR-associated changes in fluorescence show hyperpolarization of membrane potential (top) and reduction in DE rate across the soma and dendritic arbor (bottom); average of (*N* = 8 cells) with 95% CI. Note trough of membrane potential hyperpolarization is visibly delayed with respect to SWR LFP peak power (dashed vertical lines) in most cellular compartments. Due to high background levels of internal membrane-bound ASAP3 in the somatic compartment, absolute fluorescence changes recorded from the soma are smaller compared to dendrites. **d.** SWR-associated hyperpolarization from **c.** (top). Mean amplitudes (−Δ*F/F*) at peak of ripple power (pink), vs minimum in expanded SWR window (ripple peak ±100 ms; light green). Two-way ANOVA for membrane potential: *p* = 5 × 10^−5^, main effect of region; *p* = 2 × 10^−13^, main effect of SWR modulation; *F* (4, 69) = 5.55*, p* = 6 × 10^−4^ interaction of SWR×region. Maximal voltage hyperpolarization lags ripple power (*p* = 0.008 for all regions, Wilcoxon paired test). **e.** Mean time of maximal membrane potential hyperpolarization of **d.** with regard to SWR LFP peak power. **f.** SWR-associated DE rate suppression from **c.** (bottom). (*p* = 0.008 in all, Wilcoxon paired test). Two-way ANOVA for DE rate: *p* = 9 × 10^−6^, main effect of region; *p* = 7 × 10^−12^, main effect of SWR modulation; *F* (4, 69) = 0.72*, p* = 0.58 interaction of SWR×region. **g.** Average DE waveforms are significantly decreased in the basal domain inside SWRs (green) compared to outside SWRs (grey). Waveforms with 95% CI were filtered with a 5-point moving average. **h.** DE peak amplitudes are slightly decreased in the basal domain inside SWRs (*p* = 0.006) but not significantly modulated in soma or apical domains (oblique: *p* = 0.42, soma: *p* = 1.0, trunk: *p* = 0.44, tuft: *p* = 0.63, all Wilcoxon paired test). Two-way ANOVA for DE amplitude: *p* = 1 × 10^−23^, main effect of region; *p* = 0.0007, main effect of SWR modulation; *F* (4, 69) = 3.34*, p* = 0.01 interaction of SWR×region. **i.** Full-width at half-max (FWHM) of regional average DE waveforms (as in **g.**) shows significantly narrower peaks in the oblique domain (*p <* 0.0004, Wilcoxon paired test). Other regions were not significantly modulated (*p* = 1.0 for basal, soma, tuft, *p* = 0.57 for trunk). Two-way ANOVA for DE FWHM: *p* = 0.02, main effect of region; *p* = 0.058, main effect of SWR modulation; *F* (4, 158) = 1.25*, p* = 0.29 interaction of SWR×region. Unless otherwise specified, all box-and-whisker plots show segment-level Q1, Q2, Q3, and range excluding outliers; points represent individual cells; using Bonferroni correction *α* = 0.01 for *post hoc* tests performed on the per-region level, *α* = 0.05 for all other tests.

### Spatial tuning in dendrites of non-place cells

In the preceding sections, we studied the basic principles of online and offline dendritic signalling in featureless environments. We next augmented the environment with tactile and visual cues in order to study dendritic signalling during an active random foraging task on a feature-rich circular belt^43;57^. We first focused on CA1PCs which did not exhibit somatic spatial tuning (“non-place” cells). We aimed to test the following hypotheses: (1) dendrites exhibit spatial tuning; (2) dendritic segments on the same branch are likely to be similarly tuned; (3) dendritic representations of single cells span the environment (Fig 5a, individual cell example; see Fig S14 for all recorded cells). In these non-place cells, we identified that approximately 33% of dendritic segments were spatially tuned, defined as a statistically significant spatial nonuniformity of DEs (calculated using the *H* statistic^58^) compared to a shuffle distribution (Fig 5c, left). We calculated the pairwise tuning curve distances^59^ between these tuned segments, and found apparent cluster structure in this tuning (Fig 5b). Even though these were non-place cells, we asked if some low baseline of somatic spatial selectivity could be driving this structure. Although there was wide variability in spatial firing between soma and dendrites in each dendritic region (Fig 5c, right), we found that dendrites in each region were *less* coherent with the soma than chance, suggesting that the dendritic tuning we observed was not explainable by somatic backpropagation; rather, the dendritic arbor and the soma are compartmentalized to some extent. We then asked on what level this compartmentalization occurs: we found that the tuning curve distances between pairs of recorded locations on the same branch were significantly smaller than tuning curve distances between pairs on different branches, suggesting branch-level compartmentalization (Fig 5d, left). However, we found no relationship of tuning curve distance and physical distance between pairs of segments along the same branch (Fig 5d, right). The tuning of each dendrite that we observe is largely determined by its presynaptic afferents, which are fixed. However their weights may change during learning, which is the hypothesized mechanism by which somas acquire spatial selectivity^60^. We finally asked how expressive the dendritic tuning we observed was. We used a simplified linear dendrite model to assess whether the measured dendritic tuning curves of a cell, taken together, were diverse enough to generate a range of tuning curves at the soma under appropriate weights (Fig 5e; Fig S14). We found that the observed dendritic tuning curves span the space of potential Gaussian place fields much better than the somatic tuning curve alone, the somatic tuning curve with spatially delayed copies, or the same number of linearly independent random tuning curves (Fig 5e, top right), and reasonably broad tuning curves can be reconstructed with very high fidelity. These results illustrate that the dendritic arbors of non-place CA1PCs exhibit both striking internal organization and diversity of tuning, which may enable it to support arbitrary somatic place fields.

**Figure 5.**
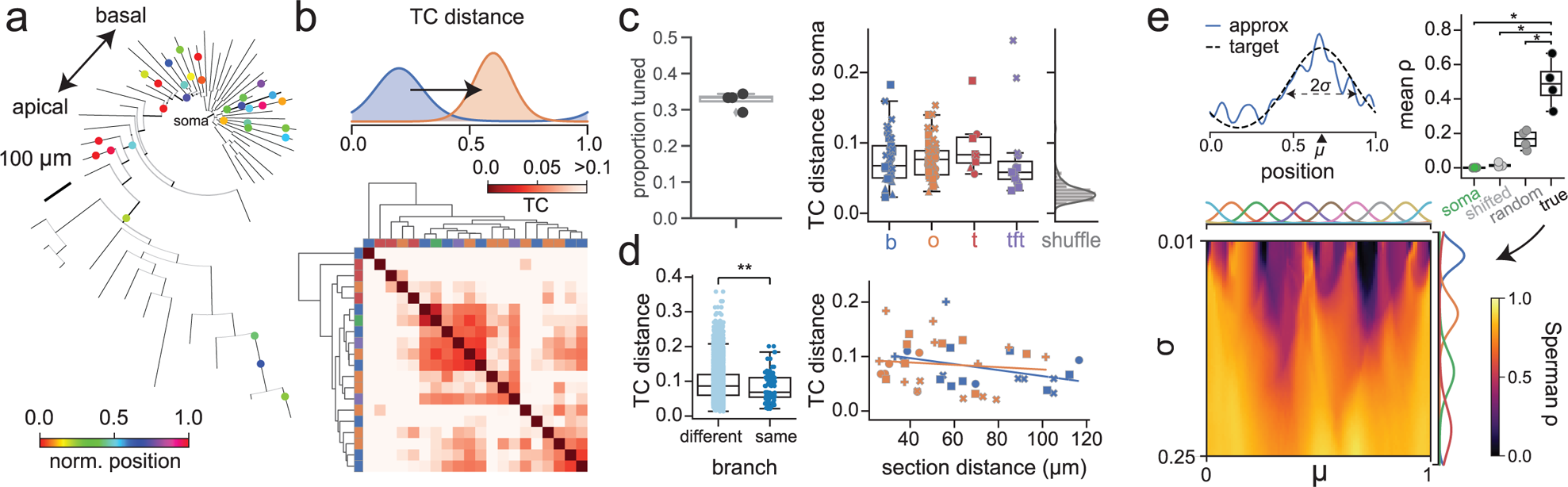
Spatial tuning in non-place pyramidal cell dendrites. **a.** Example phylogram of reconstructed cell (AN061420-1), with recording locations colored by tuning curve centroid. **b.** Example tuning curve distance (TCD, i.e. Wasserstein distance) matrix. Many segments exhibit high distance from soma. Top: Schematic of TCD calculation. TCD is normalized such that *d* = 1 corresponds to point masses on opposite sides of the belt. **c.** Left: Proportion of tuned dendrites by cell, where tuning is defined as having a p-value below the 5th percentile of a random shuffle. Right: Cotuning of tuned dendrites with their respective soma. Black: *N* = 89 tuned dendrites from 4 cells; grey: TC distance 95th percentile from shuffle performed on each dendrite. Basal: *p* = 2 × 10^−5^, oblique: *p* = 8 × 10^−9^, trunk: *p* = 0.008, tuft: *p* = 0.02, Mann-Whitney U test. **d.** Dendrites operate as compartments on the single-branch level. Left: Pairs of segments recorded on the same branch are significantly more co-tuned than pairs on different branches (*p* = 0.0009, Mann-Whitney U test). **e.** Arbitrary somatic tuning curves can be realized as sparse nonnegative combinations of tuned dendrites in non-place cells. Top left: Example tuning curve and approximation as weighted sum of dendrites. Top right: Mean realizability (Spearman’s r) of tuning curves in (bottom) using somatic tuning curve alone, somatic tuning curve with local shifts, and random tuning curves (true vs soma-only: p=0.018, true vs soma/shifted: p=0.007, true vs random: p=0.006, paired t-test). Bottom: Realizability of theoretical Gaussian tuning curves parameterized by location and width as nonnegative least-squares fit from tuned dendrites.

### Spatial tuning in dendrites of induced place cells

Finally, we tested the hypothesis suggested by the above model: that the diversity of dendritic tuning in non-place cells enables arbitrary place fields to be implanted at the soma. We carried out single-cell optogenetic place field induction (Fig 6a)^24;61^ and recorded activity from small, randomly selected sets of dendrites. We observed a significant increase in the coherence of dendritic tuning compared to non-induced cells and also found that this coherence corresponded well with the somatic peak (two example cells shown in Fig 6b,c). To quantify this observation, we computed the DE tuning vector^43^ for each dendrite and compared the diversity among tuning vectors *σ_arbor_* (defined as the circular standard deviation in the set of dendrite tuning directions) between induced and non-induced cells. Using this metric, we found that the arbors of induced cells showed significantly lower diversity, suggesting greater coherence in their dendritic representations (Fig 6d, top). Rather than binarizing tuned vs untuned, we now use a continuous metric of tuning strength (defined as the normalized magnitude of the resultant vector), and observed that among the non-induced cells, stronger somatic tuning significantly inversely correlated with diversity of tuning in the arbor (*r*^2^ = 0.74*, p* = 0.006), while induced cells exhibited low arbor tuning diversity at all soma tuning strengths (Fig 6d, bottom). Finally, we sought to dissect these differences at the level of single dendrite-soma pairs (Fig 6e). We found that somatic tuning is a poor predictor (*r*^2^ = −0.63) of dendritic tuning in general among non-induced cells, but a very good predictor (*r*^2^ = 0.95) of tuning among induced cells. We found that the variability of a single dendrite’s tuning negatively correlates to the strength of its conjugate soma tuning in both non-induced (*r*^2^ = 0.22) and induced (*r*^2^ = 0.44) cells. Interestingly, we find that place cell induction also serves to similarly organize slow plateau events in induced place cells compared to noninduced place cells (Fig S7d). Together with our previous results, these findings suggest that somatic place field acquisition reorganizes the predominant patterns of electrical activity observed at dendrites.

**Figure 6.**
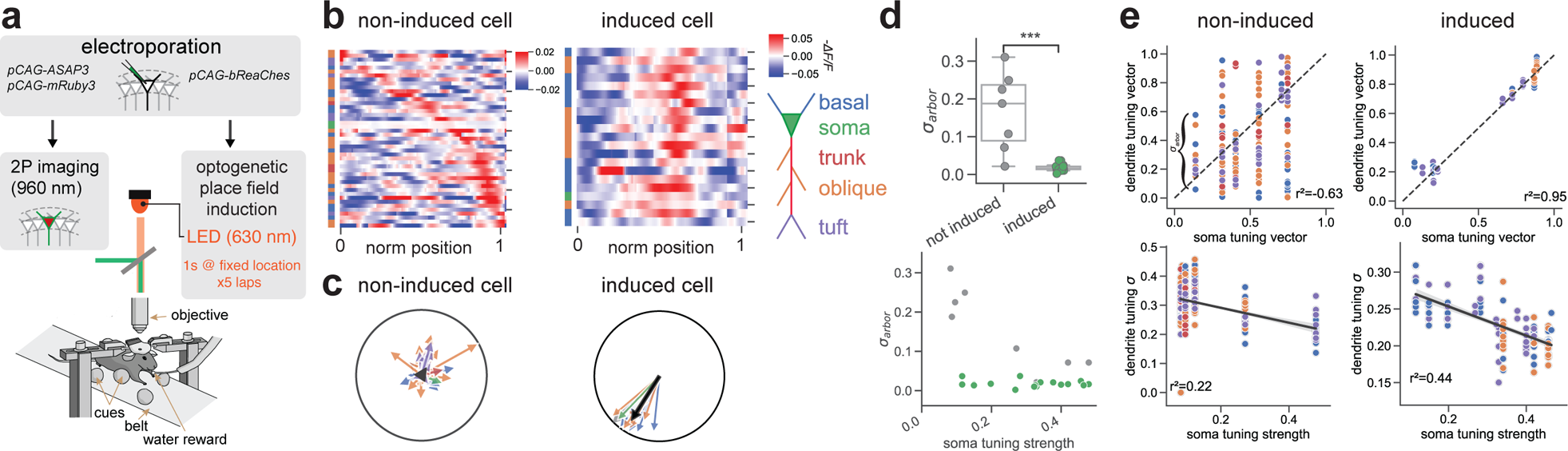
Spatial tuning in pyramidal cells dendrites following place field induction. **a.** Experimental setup. ASAP3, mRuby3 and bReaChes plasmids were single-cell electroporated in CA1 pyramidal cells. Place fields were optogenetically induced with 1s-long LED photostimulation of bReaChes at randomly chosen, fixed location on the treadmill belt for five consecutive laps. **b.** Example tuning maps of dendrites from a non-induced (non-place) and an induced (place) cell, sorted by tuning curve maximum. **c.** Mean normalized DE tuning vectors from an example non-induced (non-place) and an induced (’place’) cell. Each arrow represents a single dendrite. **d.** Tuning diversity across the dendritic arbor (*σ_arbor_*) in induced (N=15) and non-induced (N=7) cells. Top: The dendritic arbor of induced cells exhibit significantly less tuning diversity *σ_arbor_*, defined as the circular standard deviation of tuning peaks in the arbor, compared to the dendrites of induced cells (Mann-Whitney U-test, *p* = 8 × 10^−5^). Bottom: Relationship between arbor tuning diversity and strength of somatic tuning. The arbors of induced cells exhibit low diversity at all somatic tuning strengths, while a significant inverse relationship exists between tuning diversity and soma tuning strength in non-induced cells (*r*^2^ = 0.74*, p* = 0.006). **e.** Relationship between tuning parameters (tuning peak, tuning strength) at the level of individual soma-dendrite pairs in non-induced (N=7) and induced (N=15) cells. Top row: Dendrite peak locations are tightly coupled to their corresponding soma peak location (*r*^2^ = 0.95 for the line *y* = *x*, black dashed line; *N* = 123 dendrites from 15 non-induced cells) in induced cells but much less coherent (*r*^2^ = −0.63*, N* = 217 dendrites from 7 cells) in non-induced cells. Bottom row: More strongly tuned somas tend to have dendrites with more spatially coherent firing in both non-induced (*r*^2^ = −0.22) and induced (*r*^2^ = 0.44) cells, with a higher magnitude correlation for induced compared to non-induced cells.

## Discussion

The question of how anatomical structure implements computational function is fundamental to the study of neural circuits. In this work, we optically interrogated subthreshold and suprathreshold electrical activity across multiple behavior states and network oscillations in the first *in vivo*, multi-site characterization of voltage dynamics across the somato-dendritic membrane surface of principal cells in the hippocampus. Our results reveal dynamic compartmentalization of electrical signals in CA1PC dendrites, consistent with behavioral state-dependent differences in synaptic input structure and active dendritic integration^23;44^.

Our work offers new insight into the relationship between intracellular electrical signaling and population patterns across brain states. First, we find that dendrites across subcellular domains undergo prominent network state-dependent activity modulation, in which locomotion is associated with a reduced rate of depolarizing events. While previous calcium imaging and extracellular electro-physiological recordings reported an increase in firing rate of CA1PCs during locomotion^62;63^, our result is consistent with intracellular recordings showing somatic hyperpolarization and suppressed firing rate in CA1^64–66^, CA3^67;68^, and CA2^68^ PCs during locomotion, as well as with recent optical voltage recordings from CA1PCs^27^.

Second, our results answer the long-standing question of whether dendrites of CA1PCs exhibit morphologically structured oscillatory dynamics *in vivo*. The prominent gradient in intracellular theta phase along the basal-tuft axis we find during online locomotion suggests that phase-shifted extracellular theta currents carried by extrinsic inputs to CA1^37;69;70^ map onto dendrites of CA1PCs in their corresponding layer, and serve as a clock to pace suprathreshold dendritic events with a significant degree of autonomy from the soma. Our finding of opposite-sign theta phase shifts for segments in the basal and oblique domains at comparable distances from the soma suggests the sub- and suprathreshold phase shifts observed cannot simply be explained by propagation delays, but may reflect anatomical gradients in the distributions of excitatory and inhibitory inputs. The theta gradient we observe intracellularly along the basal-apical axis is reminiscent of previous extracellular LFP recordings demonstrating that theta oscillations are organized as traveling waves in CA1^71;72^. The subcellular correlates of theta oscillations we report here provide *in vivo* support for soma-dendrite or oscillatory interference models for the generation of the hippocampal temporal code during spatial navigation^33;66;73–78^.

Third, our results identify subthreshold membrane potential hyperpolarization and depolarizing event rate suppression across the somato-dendritic membrane during offline SWRs. This result is in agreement with the prominent hyperpolarization observed around SWRs with somatic intracellular electrophysiological recordings from CA1PCs *in vivo*^79–82^, (cf. ref. 83). It is important to note, however, that SWRs are associated with a relative but not absolute inhibitory state for dendrites, as DEs still occur, albeit at a lower rate, around SWRs. Sharp-wave ripples are closely associated with hippocampal replay, during which online sequences are recapitulated in the offline state^36^. Inhibition can thus play a crucial role in organizing SWR-associated replay of online activity patterns while also suppressing the reactivation of irrelevant representations^84^. The predominant dendritic inhibition in SWRs is likely brought about by GABAergic interneurons innervating CA1PC dendrites^49;85^, further implicating dendritic inhibition in structuring excitatory input integration and plasticity *in vivo*^35;61;79;86–88^. In particular, parvalbumin-expressing basket and bistratified cells, which are strongly activated during SWRs^49;85;89^ may provide inhibition to perisomatic and proximal dendritic compartments of CA1PCs during SWRs, respectively.

Finally, our voltage recordings suggest the presence of distributed feature selectivity of electrical signals in dendritic arbor in un-tuned CA1PCs, where spatially tuned branch-level compartments tiling space exhibit a high degree of within-branch coherence even in the absence of tuned somatic output. This finding is in line with recent two-photon calcium imaging studies showing functional independence and compartmentalization in CA1PC dendrites^19;24^. Recent studies also revealed that arbitrary somatic place fields can be induced in non-place cells using spatially localized electrical^34;60^ or optogenetic simulation^24;61;90;91^. Indeed, we find that following optogenetic field induction, dendritic tuning becomes significantly more coherent with somatic tuning in all compartments, likely due to backpropagating axo-somatic action potentials dominating dendritic tuning. Thus, similar to neocortical principal cells^15;21;22;92^, hippocampal place cells may also integrate distributed input-level tuning into a singular output-level receptive field through synaptic^60^ or intrinsic^93^ plasticity.

There are limitations to our study. The sequential two-photon imaging approach used here does not allow for disambiguation of backpropagating action potentials from locally generated dendritic spikes, or detailed characterization of propagation-dependent changes in event amplitude and shape. Furthermore, our study focuses primarily on single spikes, without resolving high-frequency complex spikes and plateau-burst events, which are more prevalent during place fields of place cells^34;63^. While ASAP3 improves upon previous indicators in fluorescence-voltage response as well as tolerability *in vivo*, the duration of imaging is limited by bleaching considerations, which constrained our ability to capture functional tuning profiles. Genetically encoded voltage indicators have proliferated, pushing forward optical access to neural membrane potential dynamics^29;30^. Newer-generation indicators are constantly being reported,^90;94–98^, with continual improvements to parameters such as signal-to-noise ratio, photostability, cytotoxicity, and speed. Future work may take advantage of these improvements to record across subcellular compartments for longer trials as well as longitudinally across days, which will allow an examination of how dendritic representations evolve over longer timescales. Finally, we record LFP from the contralateral hemisphere, consistent with established practice, to reduce artifacts and circumvent the difficulty of recording ipsilaterally^42;89;99^. Previous studies^47;48^ have found high coherence in interhemispheric LFP, with highly correlated theta power, only minor (< 20 deg) theta phase variance, as well as highly correlated SWR power and co-occurrence of events between hemispheres, though the coherence between hemispheres is not perfect.

In summary, we have illustrated how the dendritic arbor functionally compartmentalizes electrical signaling *in vivo*. We have shown across network and behavior states that local events are coupled to local oscillations. Our results suggest that the functional architecture of neuronal input-output transformation in CA1PCs *in vivo* is best captured as a multi-layer interaction between dendritic subunits exhibiting considerable autonomy, as previously suggested by *in vitro* and modeling studies^23;44;100–103^. This highlights the insufficiency of point-neuron models in capturing the computational capacity of either single cells or neural circuits *in vivo*. While our recording configuration did not allow for simultaneous recordings from the soma and dendrite, our findings are suggestive of an important compartmentalized component of electrical signaling, in particular during theta-associated memory encoding, which could increase dendritic computational capacity *in vivo*^8;10;104–108^. Future studies using simultaneous multi-compartmental recordings from place cells will aid in uncovering how dendritic oscillatory and spiking dynamics are utilized to implement the hippocampal time and rate code for navigation and learning^33;60;73–75;109;110^.

## Acknowledgments

We thank Dr. Jeffrey Magee for providing *in vivo* intracellular electrophysiological recordings, and Dr. Stephen S. Siegelbaum for providing *in vitro* intracellular electrophysiological recordings. We thank Drs. Liam Paninski, Kaspar Podgorski, Jeffrey Magee, and Steven Siegelbaum for their invaluable comments on the manuscript. Z.L. is supported by NIH grants F31NS120783 and T32GM007367. M.Z.L. is supported by NIH grants U01NS103464 and RF1MH114105; Post-9/11 GI Bill, and NIH grant 5T32MH020016 (S.W.E.). A.L. is supported by NIH grants 1R01MH124047, 1R01MH124867; 1U19NS104590, 1U01NS115530, and 1R01NS121106, and the Kavli Foundation.

## Author Contributions

A.L., A.N. conceived the study and wrote the manuscript; Z.L. analyzed data, wrote the manuscript; A.N. analyzed data, designed and constructed custom two-photon laser-scanning microscope with help from D.H.; A.N. developed and applied single-cell electroporation; A.N. performed surgeries together with D.M.L.; A.N. performed experiments, preprocessed and validated data, developed ASAP3 Markov model based on data contributed by G.Z. and M.Z.L; K.C.G. performed surgeries, performed experiments and preprocessed data; C.M.Y.reconstructed neuronal morphologies and contributed with illustrations; D.M.L., N.E.M. performed image analysis, segmentation, morphological registration, behavior training together with A.N.; M.C., S.W.E., J.Y., C.M., and M.Z.L. provided ASAP3 GEVI and advice.

## Lead Contact and Materials Availability

Further information and requests for resources and reagents should be directed to the corresponding authors. All unique resources generated in this study are available from the Lead Contact with a completed Materials Transfer Agreement.

## Methods

### Mice

All experiments were conducted in accordance with National Institute of Health guidelines and with the approval of the Columbia University Institutional Animal Care and Use Committee. C57Bl/6J non-transgenic mice were used for all experiments. Mice were kept in the vivarium on a reversed 12-h light–dark cycle and were housed with 3–5 mice per cage (temperature, 22–23 °C; humidity, 40%).

### Cannula implants

Imaging and single-cell electroporation cannulas, having a trapezoidal shape when viewed from the side (height 1.9 mm, top length 6.6 mm, base length 3.1 mm, width 3.0 mm), were 3D printed using 316L stainless steel (InterPRO Additive Manufacturing Group, US). Glass windows of 3.0 mm diameter and 0.13–0.16 mm thickness containing a rectangular opening of 0.2 mm by 0.35 mm offset 0.3 mm from the window center were laser cut (Potomac Photonics, US) and attached to the metal cannulas using a UV-curable adhesive (Norland optical adhesive 81, Thorlabs, US). Finally, a 0.02 mm thick silicone membrane covering the rectangular opening was glued to the glass bottom using a thin layer of UV-curable silicone adhesive (5091 Nuva-SIL, Loctite, US). This allowed glass microelectrodes to easily pierce through the membrane, repeatedly, while the brain remained insulated from the environment. Cannulas were sterilized prior to implantation.

### Surgery

Male and female C57BL/6J mice of 2–2.5 months of age were anesthetized with isoflurane and placed in a heated stereotactic mount where anesthesia was maintained during the surgical procedure. Eyes were lubricated (Puralube ophthalmic ointment, Dechra, US) and the scalp was thoroughly disinfected by three alternating wipes between 70% ethanol and povidone iodine solution. Meloxicam and bupivacaine were administered subcutaneously to minimize discomfort. The scalp was removed and the exposed skull was cleaned with 70% ethanol and dried with an air duster jet. The skin surrounding the surgery site was reattached to the skull using a tissue adhesive (Vetbond, 3M, US), while the skull surface was treated with a UV-curable adhesive (Optibond, Kerr, US) to improve dental cement adhesion.

For measurement of local field potential (LFP), a reference screw with a soldered lead was inserted to make contact with the contralateral-to-imaging cerebellar lobe, while a ground screw was similarly inserted in the ipsilateral-to-imaging cerebellar lobe. The insertion sites were dried and both screws were subsequently secured with dental cement. A small craniotomy matching the shape of the implantation cannula was made to center the imaging window at −2.3 mm AP, 1.6 mm ML. After removing the bone and dura, cortical aspiration was performed while irrigating the site with ice-cold cortex buffer to carefully expose medio-lateral axonal (ML) fibers overlying the hippocampus. These fibers could be visually separated into thicker and thinner types, the latter lying right above the anterior-posterior (AP) fibers. Thicker ML fibers were removed without disturbing AP fibers and the cannula was inserted and pushed down 1.6 mm, flush with the exposed tissue. The surgical site was dried and the cannula was initially glued to the bone using a tissue adhesive (Vetbond, 3M, US). Hippocampal LFP was measured by inserting a 0.004” outer ′ PFA-coated tungsten wire (#795500, A-M Systems, US) 1.1 mm deep from dura at same coordinates as the center of the imaging window in the contralateral-to-imaging hemisphere. In a subset of implants, the recording location of the tungsten wire was determined to be within the pyramidal layer from the shape of spontaneous ripples occurring at rest in the awake mouse. The cannula and LFP wire were fixed firmly with dental cement, which covered the skull and a headbar used for head-fixation during electroporation and imaging was attached. Ground, reference and LFP signal leads were attached to a connector strip socket (851-43-050-10-001000, Mill-Max manufacturing Corp., US) similarly fixed with dental cement. Finally, saline and buprenorphine were administered subcutaneously to minimize post-operative discomfort and the well-being of mice was monitored every 12 hours for three days. Mice were left to recover for 7 days prior to the start of treadmill habituation and the implant was left to stabilize for 3 weeks before electroporation and imaging.

### Single-cell electroporation

Endotoxin-free plasmids were purified using silica columns (NucleoBond Xtra Midi EF, Macherey-Nagel/Takara Bio, US). Plasmids expressing red fluorescent protein mRuby3^41^ at 25–100 ng/*µ*L under CAG promoter in combination with ASAP3^28^ voltage sensor at 30–100 ng/*µ*L under either CAG, EF1a or hSyn1 promoters, were electroporated in single dorsal hippocampal CA1-area pyramidal neurons under two-photon microscopy guidance using mice of <4 months of age at the time of electroporation.

Long-taper pipettes were pulled from borosilicate glass (G200-3, Warner Instruments, MA, 2.0 mm OD, 1.16 mm ID) using a horizontal puller (DMZ puller, Zeitz Instrumente Vertriebs GmbH, Germany) to yield a series resistance of 4–6 MΩ when dipped in 0.1 M PBS and filled with the following plasmid carrier saline solution (in mM): 155 K-gluconate, 10 KCl, 10 HEPES, 4 KOH, 0.166 Alexa488-hydrazide, having pH 7.3 at 25 degrees Celsius.

For the electroporation procedure, mice were placed in a heated, head-bar fixation equipped Zeiss stereo-microscope, and were anesthetized for ≈ 45 min by subcutaneous injection of a Ketamine-Xylazine mixture. The dental-cement head-cap implant, cannula, imaging window and the silicone-membrane covered opening were thoroughly cleaned with sterile 0.1 M PBS and debris was removed from the opening surface using the sharp tip of an injection needle (without piercing the membrane). A custom-designed electroporator relay unit was used to flexibly switch between a patch-clamp amplifier (BVC-700A, Dagan Corporation, MN, US) head-stage used to monitor pipette resistance and a digitally-gated stimulus isolator (ISO-Flex, A.M.P.I, Israel) used to deliver electroporating voltage pulses. Both the patch-clamp amplifier and the stimulus isolator were controlled by a digitizer (Digidata 1550B, Molecular Devices, CA, US). Pipettes containing plasmid and saline solution mix were mounted on a micromanipulator (PatchStar, Scientifica, UK) and advanced at an angle of 32 degrees through the silicone membrane under two-photon laser scanning microscopy guidance using 920 nm excitation, with 80–120 mBar positive pressure until the membrane was pierced. If pipettes pierced the membrane without becoming clogged, they were advanced several 10’s of micrometers into the tissue and the pressure was reduced promptly to 20–30 mBar to avoid spilling dye under the imaging window. More frequently, pipettes were clogged on the first piercing attempt and this was usually remedied by retracting the pipette and returning it into the saline-filled bath and applying large positive pressure alone or in combination with large voltage pulses −90 V, 100 Hz, 0.5 ms pulse ON for 1 second. After unclogging, pipettes were pushed back through the silicone membrane along the same piercing tract, and the unclogging procedure was repeated usually less than three times, as needed. After each piercing, the silicone membrane resealed and in this manner could be pierced repeatedly without losing its integrity or sterile insulation within our experimental conditions. After successfully piercing the silicone membrane, pipettes were gradually advanced toward the dorsal hippocampal CA1-area pyramidal layer, at about 120–150 *µ*m beneath the imaging window, maintaining 20–30 mBar positive pressure. Upon approaching a cell body and pipette series resistance increasing 1–2 MΩ, pressure was decreased to 6–8 mBar and a series of −4 to −5V, 100 Hz, 0.5 ms electroporating pulses were delivered for 1 second after which the pipette was kept on the cell body for an additional 4–5 seconds before slowly retracting. Successful electroporation was accompanied by dye filling the cell body and dendrites, with 4/5 cells remaining intact 10-15 minutes post SCE. Out of these intact cells, a further ≈4/5 cells expressed both plasmids 15–24h post SCE, with remaining cells usually being intact and containing residual Alexa 488. Expression of mRuby3 could be seen as early as 15h post SCE, while ASAP3 expression was evident after 24h, when Alexa488 cleared from the cell. On rare occasions only one of the plasmids was expressed. While expression levels could be varied by the choice of promoter, cell filling time, plasmid concentration and expression time, the probabilistic nature of expression introduced variability that was offset by either advancing or delaying the imaging session up to 12h. Per electroporation session, 1–2 cells were prepared in this manner, spaced 100–150 *µ*m apart to allow for unequivocal morphological reconstruction of their dendritic trees.

### Functional imaging

Head-fixed mice were habituated to run freely on a cued treadmill. Cells were imaged within 36–60h post SCE, with plasmid-expressing cells remaining healthy for ≈72h post SCE as judged by their morphology and presence of depolarizing fluorescence transients. Two-photon laser scanning microscopy was done with a custom built 8 KHz resonant galvanometer-based microscope using: a 4.0 mm 1/e^2^ ′ Gaussian beam aperture 3-galvo scanner unit (RMR Scanner, Vidriotech, US), 50.0 mm FL scan lens (SL502P2, Thorlabs, US) 200.0 mm effective FL tube lens (2x AC508-400-C Thorlabs, US, mounted in a Plössl configuration, custom large aperture fluorescence collection optics (primary dichroic: 45.0 x 65.0 x 3.0 mm T865lpxrxt Chroma, US; secondary dichroic: 52.0 x 72.0 x 3.0 mm FF560-FDi02-t3 Semrock, US; green emission filter: 50.8 mm ′ FF01-520/70 Semrock, US; red emission filter: 50.8 mm ′ FF01-650/150 Semrock, US; GaAsP PMTs for green and red channels, PMT2101 Thorlabs, US) matched to a 0.8 NA, 3 mm W.D., water-immersion Nikon 16x objective, commercial software (ScanImage, Vidriotech, US) and electronics (PXIe-1073 5-slot PXIe chassis, PXIe-7961R FlexRIO FPGA module, 5732 2-channel 80 MS/s FlexRIO DAQ, PXIe-6341 X Series DAQ, National Instruments, US). To reduce out-of-focus movement, the point spread function was axially elongated by under-filling the objective to 0.4 effective N.A. Two-photon excitation of ASAP3/mRuby3 pair was done at 960 nm (to excite mRuby3 better) using a tunable ultrafast pulsed laser source (Chameleon Ultra II, Coherent, US), modulated by a Pockels cell synced to the resonant scanning system (350-80LA modulator, 320RM driver, Conoptics, US). Transform-limited pulses were obtained at the sample using a custom SF11 glass prism-based pulse compressor.

Fluorescence scans lasting 45 seconds were acquired at ≈440 Hz (50–60x optical zoom, 32 lines/frame, 32–64 pixels/line) from preselected locations on the cell body and dendritic tree (basal, apical trunk, oblique, tuft) at various laser powers to obtain a more reproducible laser power delivery that is less dependent on the imaging depth, which bleached ASAP3 fluorescence 0.3 to 0.5 of its initial value. Under these conditions no signs of photodamage were observed (e.g. beading/swelling of dendrites), cells remained healthy throughout the recordings and morphology could be recovered.

### Single-cell optogenetic place field induction

Optogenetic place field inductions were performed as previously described^24;61^. DNA plasmid constructs (pCAGGS–ASAP3b, pCAGGS–mRuby3, pCAGGS–bReaChes) were diluted to 50ng/µL and electroporated into single dorsal hippocampal CA1-area pyramidal neurons. Successfully electroporated cells were imaged 48-72h post electroporation. Cells were stimulated for 1 second at a randomly chosen location for five laps with an ultrafast and high-power (30-40mW measured after the objective) collimated LED at 630 nm (Prizmatix, 630 nm). LED stimulation light pulses were delivered using custom-built electronics, which were synchronized with the blanking signal of the imaging beam Pockels cell, allowing pulsed LED stimulation during the flyback periods of the Y-galvanometer between each imaging frame. Following this wide-field optical stimulation protocol, the remainder of the imaging session was used to acquire postinduction somatic and dendritic data.

### Morphological reconstruction

Morphology acquisition was done using 1030 nm two-photon excitation of mRuby3, overfilling the Nikon 16x objective for 0.8 effective NA. Multiple image stacks tiling a cell were acquired at 1.0 *µ*m axial and 0.4 *µ*m transverse pixel resolution under isoflurane anesthesia. Neuronal morphologies were reconstructed from fluorescence stacks using Neurolucida (MBF Bioscience, US). Neurite thickness was set to full width half maximum (FWHM) of a fluorescent segment cross-section. Morphological path-distance measurements e.g. between imaged ROIs and the somatic compartment were done using NEURON’s^111^ inbuilt functionality using custom Python scripts.

### Phylogram rendering

We introduce the phylogenetic dendrogram, or “phylogram”, as a systematic visualization tool for dendritic morphology. This visualization is motivated both aesthetically and algorithmically by the tree of life visualization used in phylogenetics.

In brief, a tree data structure is constructed with the soma at the root (“last common ancestor” in phylogenetic terms). Intermediate nodes are added corresponding to branching points, with radial edge lengths proportional to physical distances along the dendrite and thickness optionally proportional to dendritic diameter. Leaf nodes correspond to dendritic tips. Finally, colored points are added on the phylogram corresponding to recorded locations. The tree is then laid out in polar coordinates.

The phylogram provides a schematic, programmatically generated way of visualizing dendritic morphology in 2D. Unlike a 2D projection view, the phylogram avoids ambiguities from self-intersections and faithfully represents distances. In addition to representing dendritic topology, distances can be measured directly as the radial distance from the soma to a recording location; angular distance is simply used to separate distinct branches.

Code for phylogram rendering was written using the BioPython, Matplotlib, and PlotLy packages.

### Fluorescence signal processing and event detection

Frame scans were in-plane motion corrected and fluorescence from segmented regions of interest (ROIs) was extracted using a modified version of SIMA package^112^. Rather than using an arbitrary scaled fluorescence signal, the modification allowed for converting fluorescence signals to a Poisson shot-noise equivalent number of detected photons. Since the speed of resonant galvanometers is not uniform, to maintain a faithful geometric representation of the sample, pixels across the scanfield are sampled unevenly in terms of the number of digitizer samples that are summed together. With the number of digitizer samples per pixel across the scanfield provided by ScanImage, an average fluorescence signal per digitizer sample (every 12.5 ns) per ROI patch was calculated. This value was converted to photon counts using previously calibrated shot-noise equivalent counts that were measured in Alexa-488 fluorescent-dye containing solution at varying laser powers and fluorescence signal levels. Since mRuby3 red-fluorescent protein has a small but not negligible bleedthrough in the green channel, this contribution to the green channel signal was linearly unmixed and subtracted based on a previously calibrated mRuby3 only labelled cell under same imaging conditions. To correct for bleaching, instead of fitting multi-exponential curves to ASAP3 and mRuby3 fluorescence traces, we opted for a more robust approach. Fluorescence traces were corrected by fitting and normalizing to a natural cubic spline curve having 10 knots that were geometrically spaced over 45s of recording (knots closer to each other at the start of the recording).

The modifications applied to SIMA also allowed for a frame-by-frame extraction of X- and Y-axis pixel displacement, which together with a low-pass filtered version of mRuby3 control signal (natural cubic spline with 20 ms knot intervals), were used to perform a 4th order polynomial feature multivariate regression (Scikit learn Python package^113^) and subtraction of ASAP3 movement-induced artifacts. If movement artifacts were too large, here defined as a decrease in mRuby3 fluorescence to less than 50% of its bleaching normalized baseline with a z-score more negative than −3, ASAP3 and mRuby3 fluorescence signals were blanked and excluded from analysis.

Depolarizing events (DEs) were detected using a thresholded match-filtered approach that was applied to the motion stabilized and high-pass filtered ASAP3 signal (butterworth, 4th order zero-phase, 0.2 Hz cutoff). The matched-filter template was constructed by averaging 440 Hz downsampled ASAP3 fluorescence responses obtained from the 4-state Markov model (see Methods: *ASAP3 Markov model*) in response to *in vivo* whole-cell soma recorded single action potential waveforms in mouse hippocampal CA1-area pyramidal neurons. DE match-filtered signal threshold for each recording was chosen by generating a Poisson shot-noise equivalent fluorescence signal and calculating a level that yielded a false positive rate in simulation of 0.01 Hz. To avoid subthreshold changes in the membrane potential being registered as DEs, for the match-filtered signal, a minimum 5% fluorescence-change threshold corresponding to sodium current inflection point of APs was set for dendritic recordings, while for somatic recordings, due to larger fluorescence background, this was adjusted for each cell and ranged 1%–5%. To improve DE estimation, the thresholded match-filtered signal was subjected to a peak detection algorithm that pruned DE events within a 40 ms window to keep a single largest event. This parameter was chosen as the shortest value balancing the competing objectives of temporal resolution (shorter windows better) and reliability (longer windows better), and cross-validated across different sizes, with good agreement apart from very short (sub-10 sample) window sizes (Fig S5). DEs were considered to be isolated if 45 ms before or after the event there were no other events.

### ASAP3 response characterization

A comprehensive characterization of the biophysical and electrophysiological properties of ASAP3 *in vitro* and *in vivo* is reported in^28^. We use HEK293 cells for calibration of the voltage-fluorescence response: given a ground-truth voltage waveform, we sought to measure the corresponding fluorescence signal and use it to both construct a fluorescence template for AP-like events as well as to calibrate a voltage-fluorescence response model. ASAP3 construct was cloned in pcDNA3.1/Puro-CAG plasmid backbone and used to transfect HEK293 cells with Lipofectamine 3000 (ThermoFisher, US) using 400 ng DNA, 0.8 *µ*L P3000 reagent and 0.8 *µ*L Lipofectamine. Cells were patch-clamped at 22–23 ^°^C and at 37 ^°^C after 24h from transfection using a Multiclamp 700B amplifier (Molecular Devices, US) and either voltage stepped for 1s starting from a holding potential of −70 mV to voltages between −200 and +120 mV and back or voltage-clamped using *in vivo* somatic measured mouse hippocampal CA1-area pyramidal neuron action-potential burst waveforms. Simultaneous with voltage stepping, ASAP3 fluorescence was excited using UHP-Mic-LED-460 LED (Prizmatix, US) through a 484/15-nm excitation filter and cells were imaged using a fast iXon 860 EMCCD camera (Oxford Instruments, US) at a frame rate of 2.5 KHz. Steady state ASAP3 fluorescence responses were described well by sigmoidal functions (Table S1), while dynamic step responses could be fit well to single and double exponential kinetics (Table S2).

The use of voltage clamp with a ground-truth AP waveform reduces dependence on the electro-physiological properties of the cell used for calibration; the fluorescence response in culture will be comparable to the fluorescence response *in vivo* assuming only that the dynamics of the sensor are comparable and that the sensor is expressed similarly in both preparations.

### ASAP3 Markov model

A minimal 4-state linear chain Markov gating model formalism was used that was similar to previous descriptions of ion-channel gating^114^, except that in the case of ASAP3, instead of modelling current conduction, the ensemble fluorescence brightness was modelled. To describe voltage-dependent transition rates between any two states, a linear thermodynamic model was used:

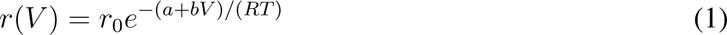

where *r*_0_, *a* and *b* are transition-dependent constants, *V* is the membrane potential, *R* is the universal gas constant and *T* is temperature. The 4-state linear-chain voltage-dependent Markov model transition matrix was then:

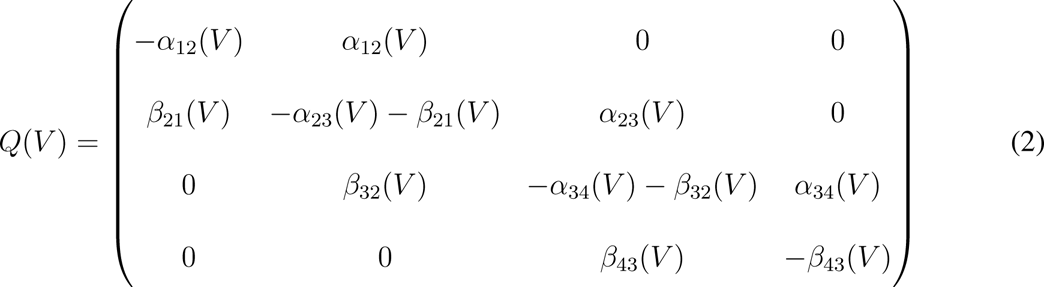

where *α_ij_*(*V*) and *β_ij_*(*V*) are voltage-dependent transition rates from state *i* to *j* and back respectively. The temporal evolution of the state probability vector *π* is given by:

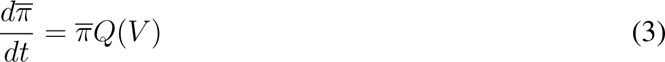

which can be numerically calculated at each time-step Δ*t*:

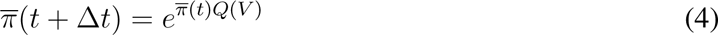

To obtain the steady state probability vector *π_∞_*, the following equations were solved algebraically:

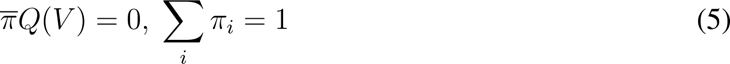

Time-dependent ensemble fluorescence for state brightness vector *λ* was calculated as:

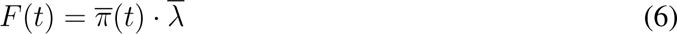

The 4-state linear-chain Markov model parameters were fit to the steady state and voltage step responses of ASAP3 in voltage-clamped HEK293 cells at 22–23 ^°^C and 37 ^°^C (Tables S1, S2) using boundary constrained limited-memory BFGS algorithm (L-BFGS-B) implemented by the Python SciPy optimization package.

### Simulation of domain-specific average DE waveforms

To obtain average domain-specific DE waveforms as shown in (Fig 1e), in response to somatically recorded APs *in vivo*, single APs were isolated from adult mouse dorsal hippocampal CA1PCs whole-cell current-clamp recordings that were kindly provided by Dr. Jeffrey C. Magee (Baylor CM, TX) from ref. 60 (baseline=-58.2±0.6 mV, amplitude=93.1±1.0 mV, FWHM=1.2±0.0 ms, mean±SEM). AP waveforms sampled at 20 KHz were converted to fluorescence responses using the above ASAP3 Markov model and downsampled to 440 Hz to match the imaging frame rate. Poisson shot-noise limited fluorescence waveforms corresponding to single isolated APs were generated by randomly picking DE fluorescence baseline photon counts from the distribution of measured DE baselines corresponding to each domain and applying a minimum 100 cts/sample threshold. Matched-filter event detection and averaging was done similar to the measured fluorescence waveforms.

### Local field potential (LFP) measurement and preprocessing

Hippocampal LFP was measured by inserting a shielded 18-pin electrode adapter board for headstages (C3418, Intan Technologies, US) in the surgically-fixed connector strip carrying LFP signal, ground and reference. To this adapter board, an RHD2132 16-channel amplifier with ground and reference pins disconnected from each other (C3334, Intan Technologies) was attached. LFP measurement was done using an RHD2000 interface board (C3100, Intan Technologies, US) that simultaneously recorded laser-scanning frame trigger and treadmill synchronization signals at a sampling rate of 10 KHz. LFP and treadmill position signals were resampled offline to about 440 Hz (and in a few cases 220 Hz) in sync with the acquired frames.

### Behavioral state classification

Behavior of mice was defined by their locomotion activity on a treadmill with tactile cues. The 2-m long cue-rich fabric belt was constructed by stitching together 3 fabrics and then adhering local tactile cues (e.g., velcro, glue gun spikes, silver glitter masking tape, green pom poms). If movement velocity dropped below a threshold of 2 mm/s for at least 1 second, mice were considered at rest. Walking/running epochs were defined to have a sustained velocity above 10 mm/s for at least 1 second.

### DE rate estimation

DE rate is estimated by counting the total number of events in a behavior state (running or nonrunning) and dividing by the duration in that state. For the purposes of rate estimation, all scans from a single segment are concatenated and only epochs of at least 1 second in duration are included.

### Analysis of theta-band dendritic fluorescence oscillations

Theta-band LFP phase was determined by first applying a zero-phase 20th total order 5–10 Hz Butterworth filter, assigning a phase of 0 deg to oscillation troughs and interpolating the phase between adjacent troughs using a linear function. Fluorescence recordings were split according to the animal’s behavior state to be either still or walking/running. To avoid bias from soma-modulated back-propagating action potentials, fluorescence samples from matched-filter detected depolarizing events were excluded within a 20 ms window centered on each event.

This process allows us to assign a theta phase to each recording sample during locomotion based on the theta-band LFP oscillation. The amplitude *A* and phase *ϕ* of the membrane potential oscillation relative to the extracellular oscillation were estimated using the discrete Fourier transform (DFT) estimator of sinusoidal parameters under additive white Gaussian noise (AWGN)^115;116^

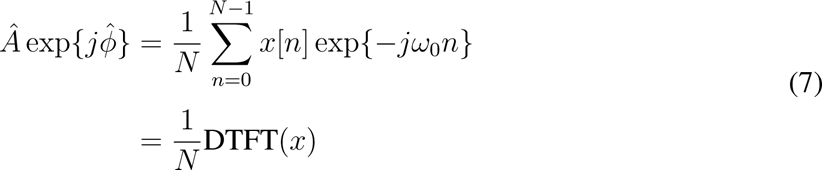

The variance of the amplitude and phase estimates derived in this way are well-characterized

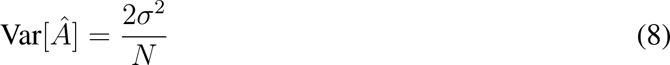

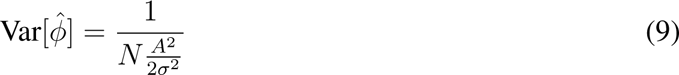

where *σ*^2^ is the noise variance, estimated empirically as the residual variance.

Segments whose amplitude estimate was significantly greater than 0 (i.e., 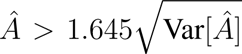) were considered to be significantly phase modulated in *V_m_*. The advantage of this estimator is its ability to detect modulation at any constant phase offset. If a significant modulation is detected, the phase offset (relative to the contralateral theta) is read off from the estimator. As variability in the placement of the contralateral LFP electrode could also introduce a constant phase offset, we report all phases relative to the estimated phase at the somatic/proximal basal compartment.

DE phase preference was calculated by first assigning each DE a unit modulus complex number exp*{jϕ}*, where *ϕ* is the phase of extracellular theta LFP oscillation at which the event occurred. The mean of these complex numbers was computed for each recording location: the argument corresponds to the estimate of the DE preferred phase, while the modulus (elsewhere called mean resultant length, MRL) corresponds to the degree of preference. We note that this method is equivalent to the DFT method if samples are observed at uniform intervals. Segments for which *p <* 0.05 under a Rayleigh test of DE phases were considered significantly phase-modulated in DEs.

### Detection of hippocampal ripples

Ripple detection was done by: 1) applying a zero-phase 4th total order (2nd order forward and backward) 80–220 Hz Butterworth band-pass filter to LFP measured during behaviorally still periods, 2) squaring the filtered signal to calculate the instantaneous power and smoothing it using a zero-phase 4th total order 25 Hz cutoff low-pass Butterworth filter, 3) scoring the instantaneous power using Tukey’s inter-quartile range (IQR) and thresholding ripple epochs at an IQR level above 10, 4) expanding ripple epochs in both directions until the IQR score decreased below 2, 5) ignoring ripple events shorter than 15 ms and joining ripple events that are less than 15 ms apart.

### Phase gradient estimation

Small variations in electrode placement could introduce random offsets to the extracellular theta phase; to mitigate this, we report all phases relative to the somatic phase, inferred by performing linear regression of the phases of significantly phase-modulated basal dendrites vs path distance for each cell; if the resulting regression is significant, we use the estimated intercept, otherwise, we take the mean phase. The relative phase determined in this way is defined as 0 and all subsequent phase estimates are reported relative to this inferred somatic phase. We show that the phase estimated this way agrees well with the true somatic phase for cells for which both were recorded.

We assessed the presence of a phase gradient in the membrane potential oscillation along the basaltuft axis both parametrically and nonparametrically. Throughout, we use a signed soma path distance convention, i.e. basal ROIs distance is negative. To estimate the phase gradient parametrically, we performed least-squares linear regression of relative phase of significantly phase-modulated dendritic segments vs path distance to soma. To estimate the same gradient nonparametrically, we binned a complete theta cycle into 20 phase bins and performed rank-regression, regressing the rank of each segment’s soma path distance vs the phase bin corresponding to that segment’s minimum membrane potential. Pearson correlation coefficients and p-values were calculated in both instances.

Recording locations in the oblique and trunk domains have two quasi-orthogonal contributions to their soma path distance: the distance from the soma at which obliques branch off the trunk axis (axial distance) and the distance of the recording location to this origination point on the trunk (radial distance). To interrogate the relative contributions of these components to the phase gradient, the phase variable was regressed simultaneously against these two distances using the *statsmodels* Python package regression.linear model.OLS function, employing an ordinary least squares method. Phase gradients within said dendritic domains were deemed to be present if the p-value associated with the regression was below 0.05.

Basal-oblique and basal-tuft dendritic domains phase span was measured as the phase difference between basal and oblique or basal and tuft dendritic domains by using either the mean phase of all recording locations within the domain, if the phase-distance regression was not significant, or the phase associated with the furthest recording location from the soma according to the regression model, if the regression model was significant.

### Phase modulation of DE amplitude

In order to ask the question of whether DE amplitude was modulated by phase in the theta cycle, we performed linear-circular fits to the empirical histograms of amplitudes and phases (as phase is a circular variable). This translates to solving the least-squares problem:

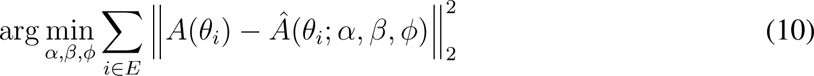

where *E* represents the set of events, *A*(*θ_i_*) is the empirical amplitude of event *i* occurring at phase *θ_i_*, and 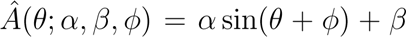 for amplitude *α*, phase *ϕ* and intercept *β*. Fitting and significance assessment were performed using the `statsmodels` package in Python.

### Peri-ripple depolarizing event rate

Significantly rate-modulated segments (Fig 4a, S11a), were defined as segments which differed significantly in Poisson rate inside and outside SWRs. The following standardized Poisson rate-difference statistic of^117^

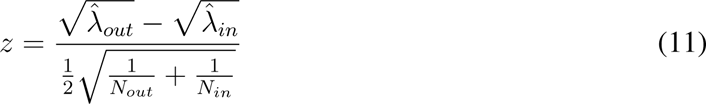

was computed, where

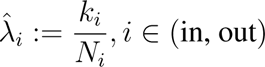

i.e., the total number of events observed in a condition *k_i_* divided by the total number of samples from that condition *N_i_*. Significance was determined by a two-sided *z*-test at a significance level of *α* = 0.05. For subsequent analyses, all segments were used without pre-filtering for segment-level significance.

### Peri-ripple depolarizing event waveforms and amplitudes

Depolarizing event waveforms inside and outside the LFP ripple epoch were calculated as an average of a 5-point moving average −Δ*F/F* in a ±100 ms window aligned to the DE event time, as determined by the previously described matched filtering approach. Event amplitudes were taken as the −Δ*F/F* value at the DE event time.

### Peri-ripple membrane potential analysis

As LFP ripple events are brief, the membrane potential trough may actually be reached outside the ripple itself. The peri-ripple *V_m_* trough was identified on a per-segment level as the minimum membrane potential value in the average segment −Δ*F/F* waveform computed in an expanded window of the LFP SWR power peak ±200 ms.

### Spatial tuning analysis

Spatial tuning analyses were performed on detected depolarizing events. Tuning curves were calculated by binning the track into 100 place bins and computing the average DE rate in locomotion epochs within each bin.

Dendritic spatial tuning may be single- or multi-peaked, driven by multiple distinct spatially-tuned inputs converging onto the same stretch of dendrite. We called a segment “spatially tuned” if its DE rate tuning curve differed significantly from shuffle. We took the *H* statistic of nonuniformity described in^58^ as our base tuning metric, and discarded segments which were not significantly nonuniformly tuned to *p <* 0.05 under this metric. A null distribution on this statistic was constructed by randomly permuting DE times within locomotion epochs 10,000 times for each segment and calculating the *H* statistic on each permutation, and the 95th percentile of this distribution was chosen as the critical value for determining significance.

### Circular Wasserstein distance

Distance between tuning curves (TCD) was computed using the circular Wasserstein distance, defined as in eq. 10 of ref. 59:

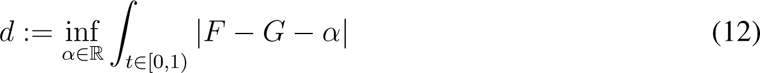

where *F* and *G* are the respective cumulative density functions (normalized to unit norm), and *t ∈* [0, 1) corresponds to normalized position around the circle.

The circular Wasserstein distance (also known as earthmover’s distance or Monge-Kantorovich distance) measures the total “cost” associated with transforming one distribution into another in terms of transporting probability mass. For single-peaked distributions, this corresponds to the usual intuition of moving the “mass” of one peak to another: the further apart the peaks, the higher the cost. Unlike peak-based metrics, the Wasserstein distance is robust to multipeaked distributions.

### Construction of somatic tuning curve from dendritic tuning curves

The place-cell somatic tuning curve is classically modeled as a Gaussian distribution on the circle, parameterized by an angular mean and variance. The tuning curves of spatially-tuned dendrites were taken as the basis set, and we asked whether any somatic tuning curve could be approximated up to scale as a nonnegative linear combination of these basis variables. We generated circular Gaussian tuning curves for a range of *µ* and *σ* parameters and computed the mean reconstruction quality as the average Spearman’s *ρ* between the NNLS reconstruction and the idealized tuning curve. We compared several alternative hypotheses: the soma tuning curve only, spatially shifted copies of the soma tuning curve, and the same number of linearly independent random tuning curves as significant dendrites.

### Tuning resultant vector analysis

We used tuning resultant vector analysis following^43^ to quantify how dendritic event tuning changed with place field induction. Briefly, every DE during locomotion was assigned a complex number corresponding to the position in which it occurred, and the resultant vector was computed as the complex mean of all of these vectors, normalized in magnitude from [0,1]. The argument (direction) of this resultant corresponds to the direction of tuning, while the magnitude corresponds to a continuous measure of the strength of tuning. Tuning variability was defined within a dendrite as the circular standard deviation of the events that occurred within that dendrite. Tuning diversity was defined across the arbor as the circular standard deviation of the resultant tuning *vectors* observed at all dendrites across the arbor.

## Data and Software Availability

*Source data are provided for each Main and Extended Data Figures. The data analyzed for this study are available at XXX. The codes used for this study are available at XXX. (*upon publication).

## Supplementary information

**Figure S1.**
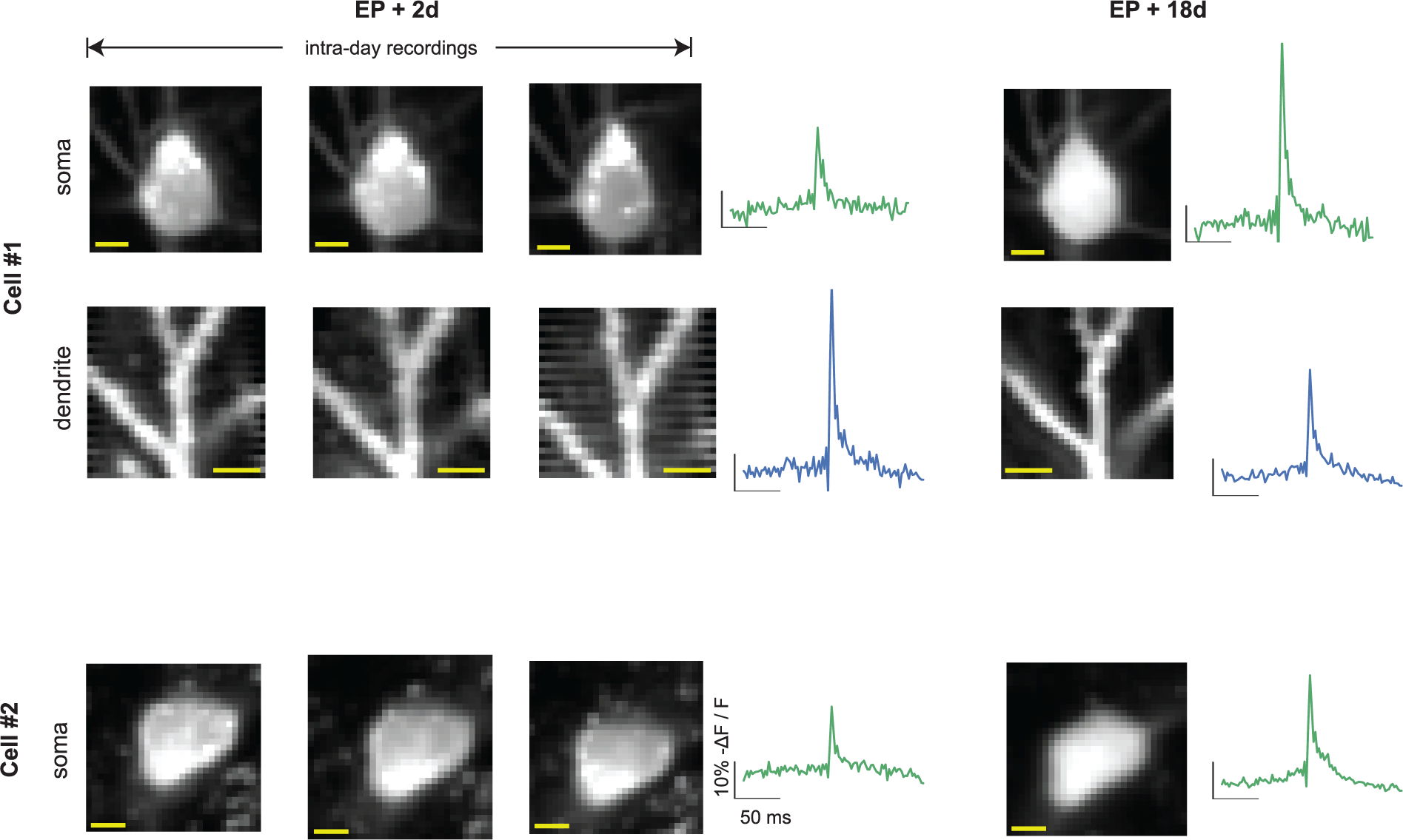
Cell health over time. Two example cells shown. We return to the same cell and same dendrites 2 days post-EP and 18 days post-EP and in recordings taken at multiple timepoints find the cell membrane intact with absence of noticeable apoptotic blebbing (scale bars 5*µ*m). The mean detected DE at each ROI over all sessions is shown in the early and late recordings: waveform morphology remains stable although amplitude may change due to changes in sensor expression.

**Figure S2.**
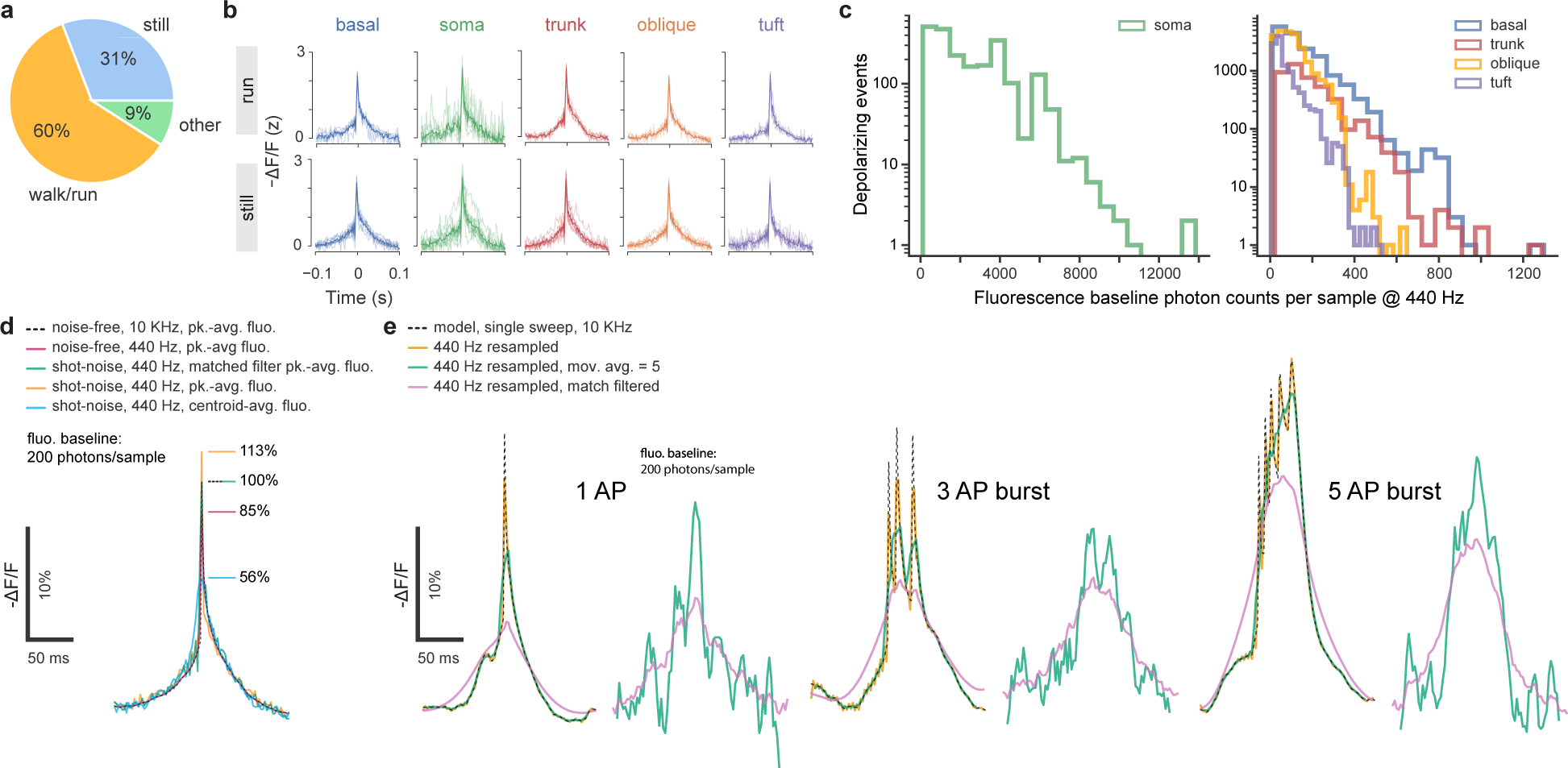
Extended data associated with Fig 1. **a.** Percent of recording time classified as locomotion, stillness, and other (transition between stillness and locomotion). **b.** Average DE waveforms (z-score) during running and stillness per dendritic region for DEs at a fluorescence baseline SNR >10. **c.** Photon counts for fluorescence baseline at time of depolarizing events (DEs) in soma (left) and dendritic domains (right) at a sampling rate of 440 Hz. **d.** Effect of downsampling to 440 Hz and choosing different methods of aligning ASAP3 Markov-model fluorescence response peaks to multiple in vivo somatically-recorded single action potentials from a mouse dorsal CA1-area hippocampal pyramidal neuron under Poisson shot-noise (200 cts/sample). Downsampling alone to 440 Hz (purple) reduced the noise-free peak-averaged response to 85% of its original value sampled at 10 KHz (black dashed). Aligning shot-noise limited waveforms by the fluorescence peaks artificially biases the peak of the average waveform (orange) to 133% of its downsampled value, while aligning waveforms using the matched-filtered signal peaks (green), the bias reduced to 118% of its downsampled value. Aligning by the centroid (cyan) of shot-noise limited waveforms severely reduced the measured peak to 66% of its downsampled value. **e.** Effect of applying a 5-point moving average filter or a matched filter to a shot-noise limited (200 cts/sample) 440 Hz recorded ASAP3 Markov-model fluorescence response to one, three and five action-potential burst waveforms. For the matched filter template, the fluorescence response to a single somatically-recorded action potential was used. Note that from the matched-filter signal it is not possible to distinguish the number or timing of action-potentials in a burst.

**Figure S3.**
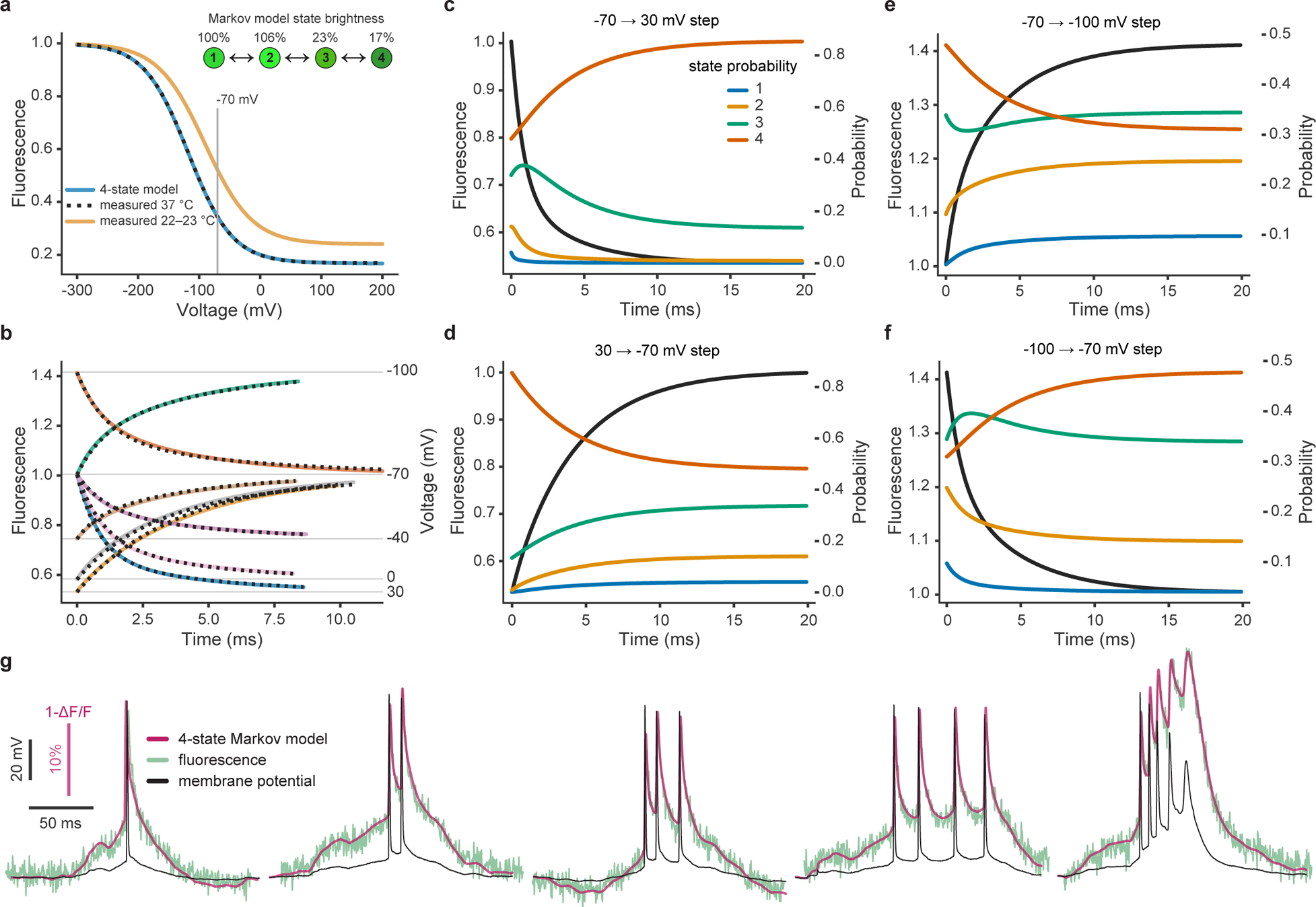
ASAP3 Markov model. **a.** Comparison of measured steady state fluorescence response at 22–23 °C (orange) and 37 °C (black dashed) and Markov model response (light blue) in voltage-clamped HEK293 cells. Note that steady state responses are temperature sensitive. Inset top: 4-state Markov chain with relative state brightness. **b.** Comparison of measured fluorescence response (black dashed) to step changes in holding potential in HEK293 cells at 37 °C and 4-state Markov model response (continuous curves). **c.–f.** Dynamic distribution of Markov model state probabilities as a function of voltage step in HEK293 cells at 37 °C. **g.** Markov model response (purple) to in-vivo measured action-potential burst waveforms (black) from a mouse dorsal hippocampal CA1-area pyramidal cell and measured fluorescence response (green) in a voltage-clamped HEK293 cell at 37 °C.

**Figure S4.**
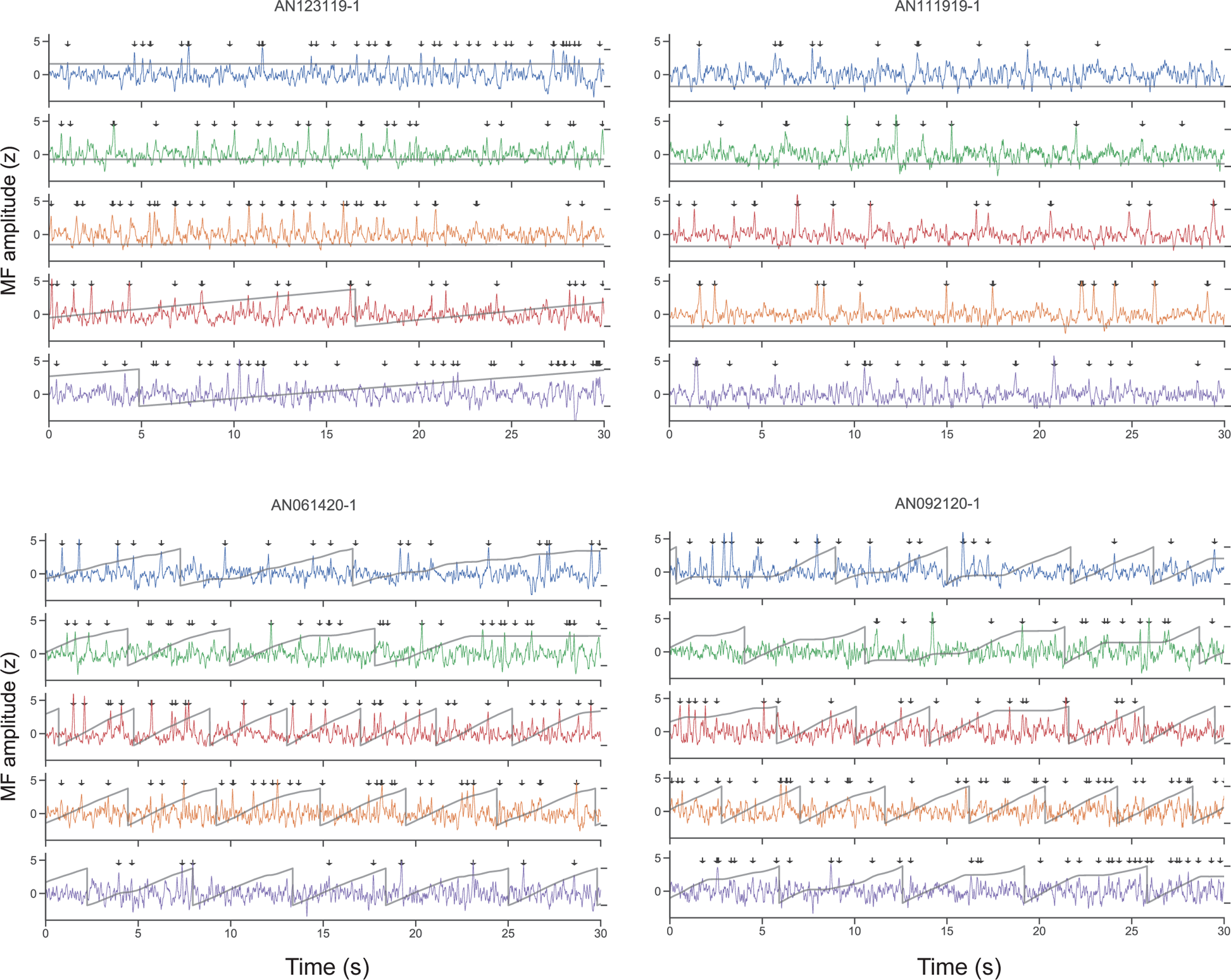
Further examples of raw signals from four distinct cells and compartments. As in Fig 1, signals, sampled at 440 Hz, were stabilized against movement artifacts, match-filtered (MF) and thresholded for depolarizing event (DE) detection (arrows) at a false positive rate of at most 0.01 Hz, plotted as −ΔF/F to reflect the conventional direction of change in the membrane potential. Grey traces: animal position during the recording.

**Figure S5.**
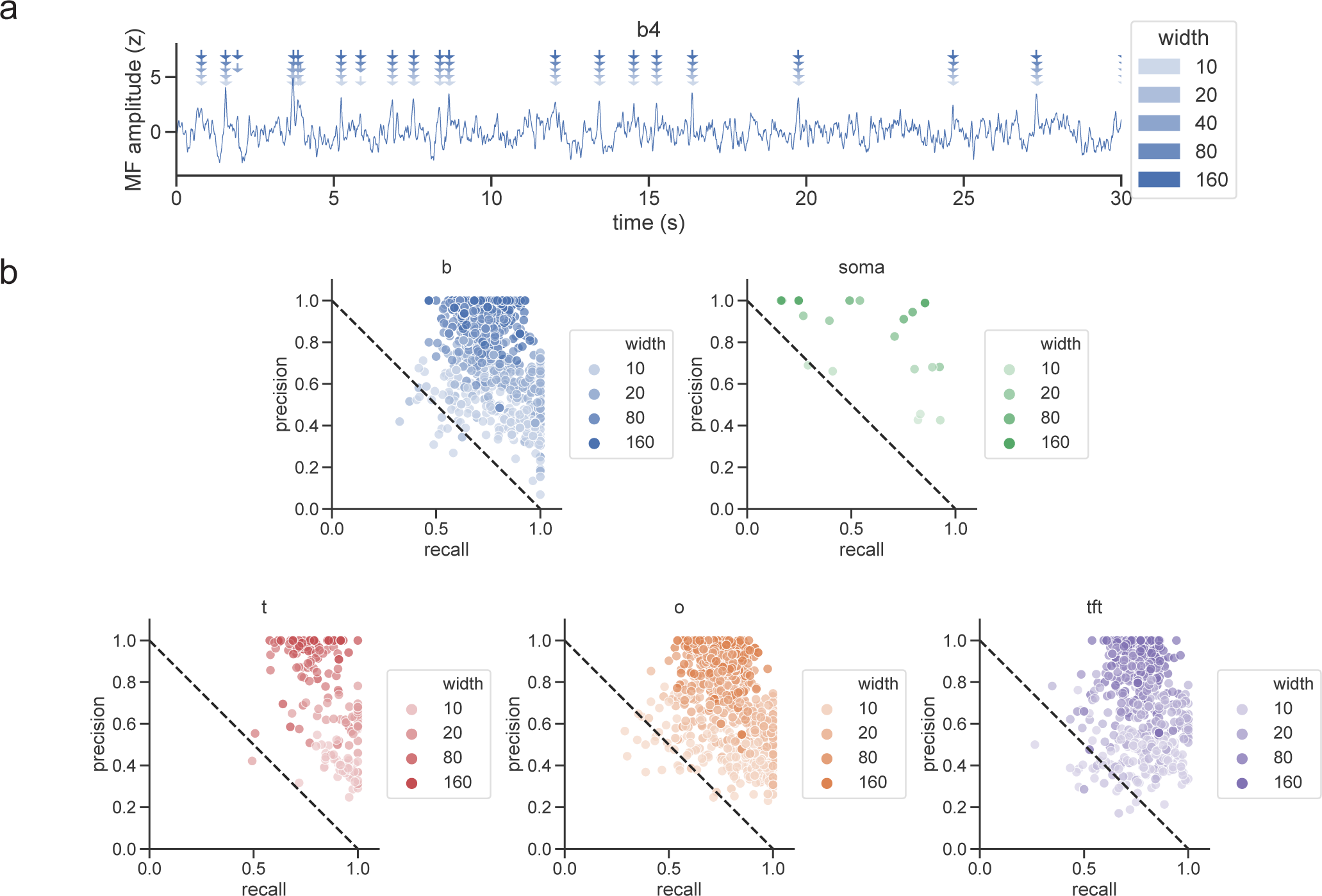
Detected events are robust to choice of event detection window. **a.** Example trace from a basal dendrite with events detected using different windows (arrows; 10 ms, 20 ms, 40 ms (default), 80 ms, 160 ms). While some variability is seen, there is agreement across all window sizes on the vast majority of events, particularly high-amplitude events. **b.** Precision-recall scatterplot by region for different choices of window size, calculated relative to the default window size (40 ms). Each point represents all events from a single dendrite. Precision is defined as # events on which the default and test window agree / total # events detected using the test window size. Recall is defined as # events on which the default and test window agree / total # events detected using the default window size. In all regions, points are concentrated in the top right in each plot indicating good agreement in detected events between different window sizes.

**Figure S6.**
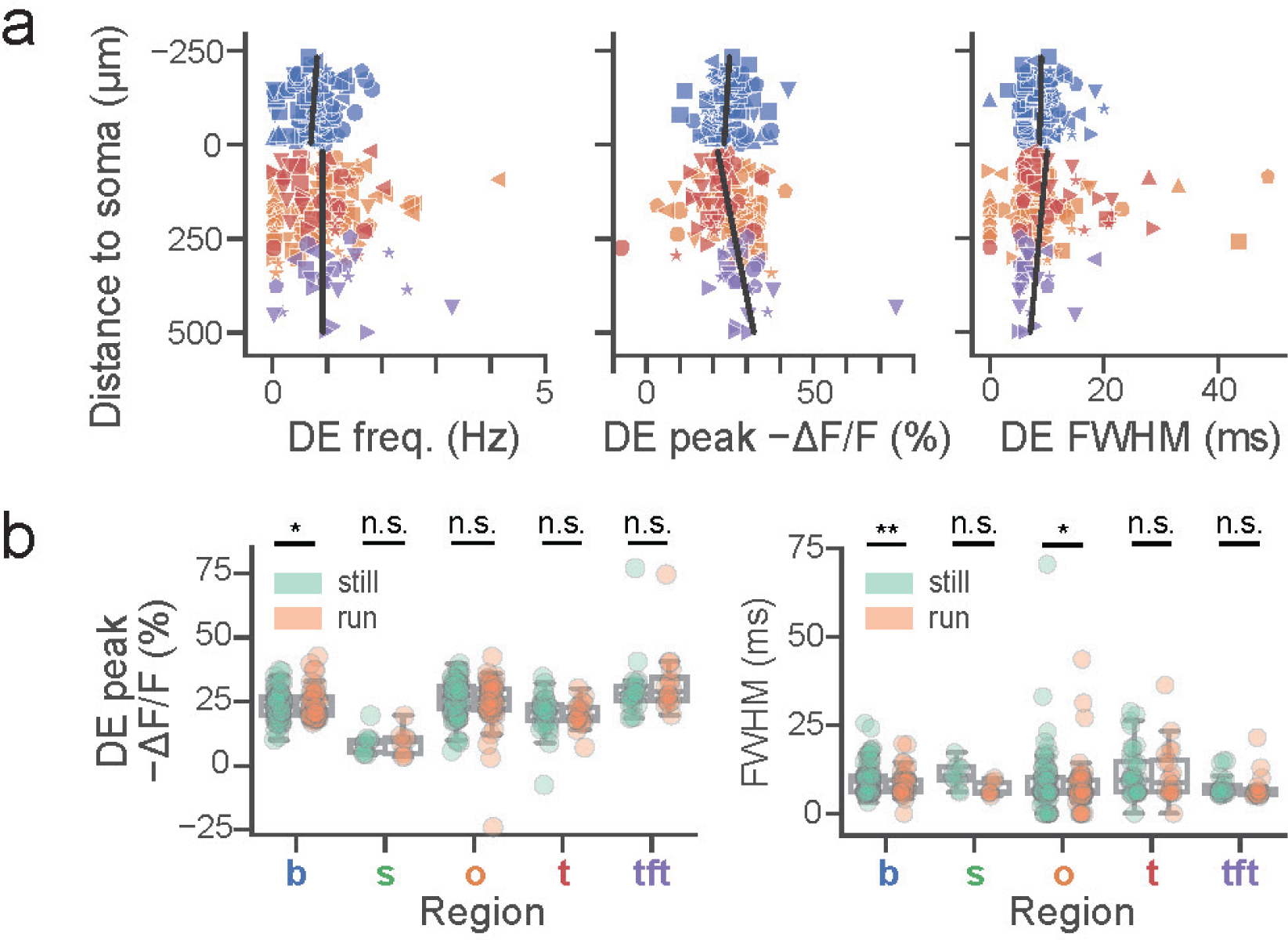
Basic characterization of DE parameters by domain and distance to soma. **a.** DE frequency (basal: slope=5 × 10^−4^ Hz/*µ*m, *p* = 0.43, *r* = 0.08; apical: slope=5 × 10^−6^ Hz/*µ*m, *p* = 0.99, *r* = 7 × 10^−4^), amplitude (basal: slope=7 × 10^−3^ %Δ*F/F/µ*m, *p* = 0.43*, r* = 0.08; apical: slope=0.02 %Δ*F/F/µ*m, *p* = 2 × 10^−4^*, r* = 0.29) and FWHM (basal: slope=1 × 10^−3^ ms/*µ*m, *p* = 0.79*, r* = 0.03; apical: slope=−6 × 10^−3^ ms/*µ*m, *p* = 0.25*, r* = −0.09) dependence on path-distance from soma across dendritic domains (same color coding) for fluorescence baseline SNR >10 (*N* = 10 cells). **b.** Left: DE amplitude is not significantly modulated between locomotion and stillness for soma (*p* = 0.55), trunk (*p* = 0.06, *N* = 1 − 12 ROIs per cell), oblique (*p* = 0.43, *N* = 6 − 40 ROIs per cell), and tuft (*p* = 0.71, *N* = 1 − 11 ROIs per cell) regions for fluorescence baseline SNR >10, *N* = 10 cells. A small but statistically significant locomotion-associated increase in amplitude was observed in the basal domain (*p* = 0.01; *N* = 5 − 30 ROIs per cell, all Wilcoxon paired test). Two-way ANOVA for amplitude: *p* = 6 × 10^−25^, main effect of region; *p* = 0.92, main effect of locomotion; *F* (4, 401) = 0.63*, p* = 0.64 region×locomotion interaction. Right: DE full-width at half-maximum (FWHM) is slightly shortened during locomotion in the basal (*p* = 1.5 × 10^−3^) and oblique (*p* = 1.2 × 10^−2^) domains. FWHM is not significantly modulated in the other domains (soma: *p* = 0.11, trunk: *p* = 0.33, tuft: *p* = 0.24; all Wilcoxon paired test). Two-way ANOVA for FWHM: *p* = 0.01, main effect of region; *p* = 0.23, main effect of locomotion; *F* (4, 401) = 0.39*, p* = 0.82 region×locomotion interaction.

**Figure S7.**
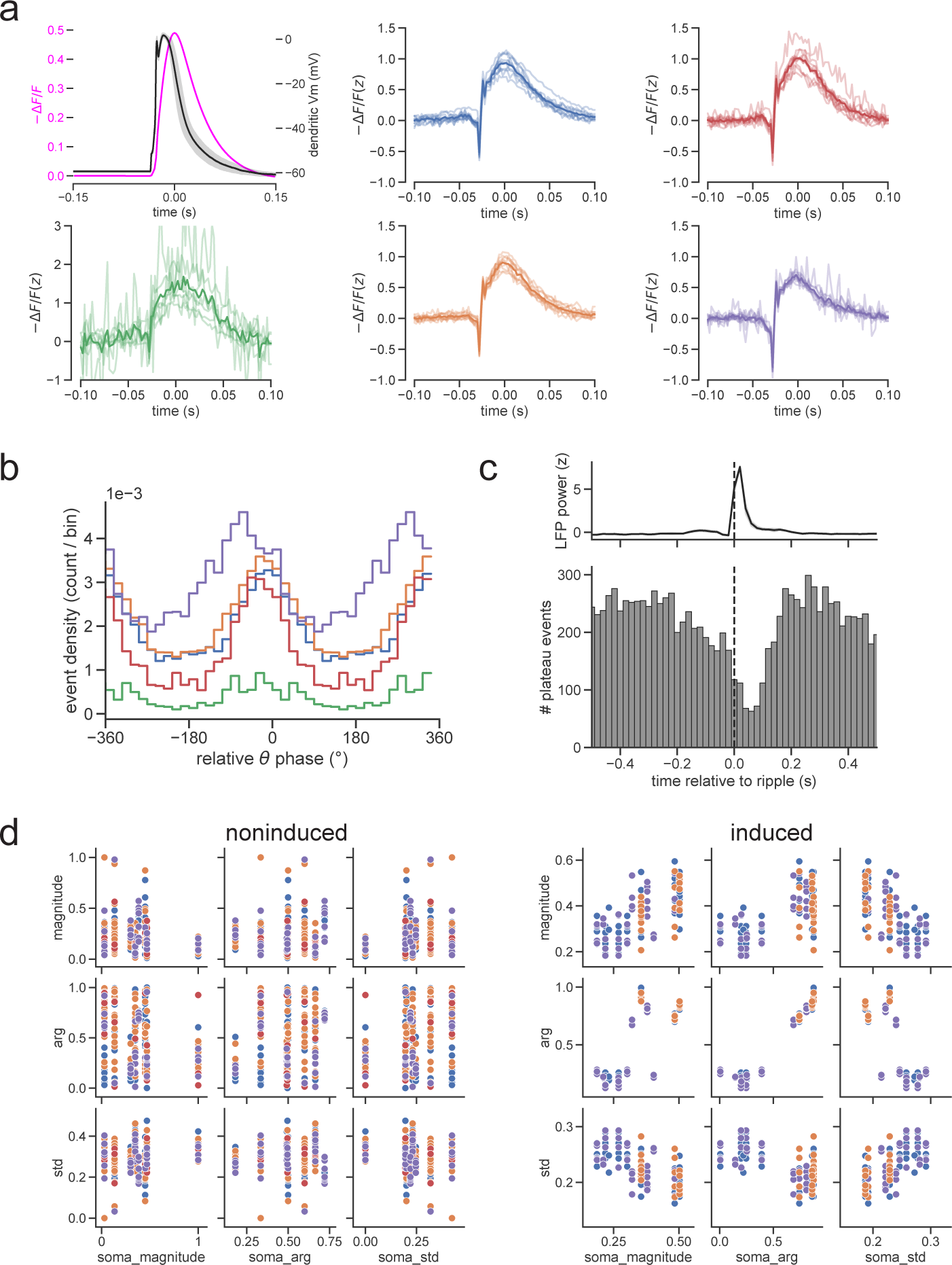
Basic properties of “slow” dendritic events calibrated to patch data. Slow events exhibit a qualitatively different waveform from fast (AP-like) events, with similarities and differences in other properties. **a.** Matched-filter based slow event detection from N=9 cells. Top left: Electrophysiologically recorded slow dendritic event waveform from ^46^ (black line; stimulation artifact removed), and model-predicted fluorescence waveform (magenta line). Colored panels: Mean slow event waveform detected in each region. Light traces: mean event from 1 cell and region. Dark trace: mean event across cells. **b.** Histogram of slow event phases by region. We note that, unlike fast events which exhibit marked phase advancement across regions (Fig 2), the distribution of slow events appears to be concentrated around 0°in all regions but the tuft. **c.** Peri-ripple slow event histogram. Dashed line: peak of LFP ripple power as in Fig 4b. Similar to fast DEs (Fig 4), slow events are suppressed around SWRs with the onset of suppression preceding the rise in ripple power, the nadir coming slightly after the ripple power peak, and a rapid return to baseline. **d.** Soma vs dendrite tuning vector relationships pre- and post place field induction. Magnitude refers to the strength of event tuning. “Arg” refers to the tuning center of mass around the circle (normalized such that 1 refers to 2*π*). “Std” refers to standard deviation. In the pre/non-induced dataset, there is no relationship between somatic and dendritic parameters, while after induction, consistent with our DE findings, dendrites’ tuning centers shift towards their somatic tuning center; strength of somatic tuning correlates to strength of dendritic tuning and inversely to variability in dendritic tuning. Each point: 1 dendrite.

**Figure S8.**
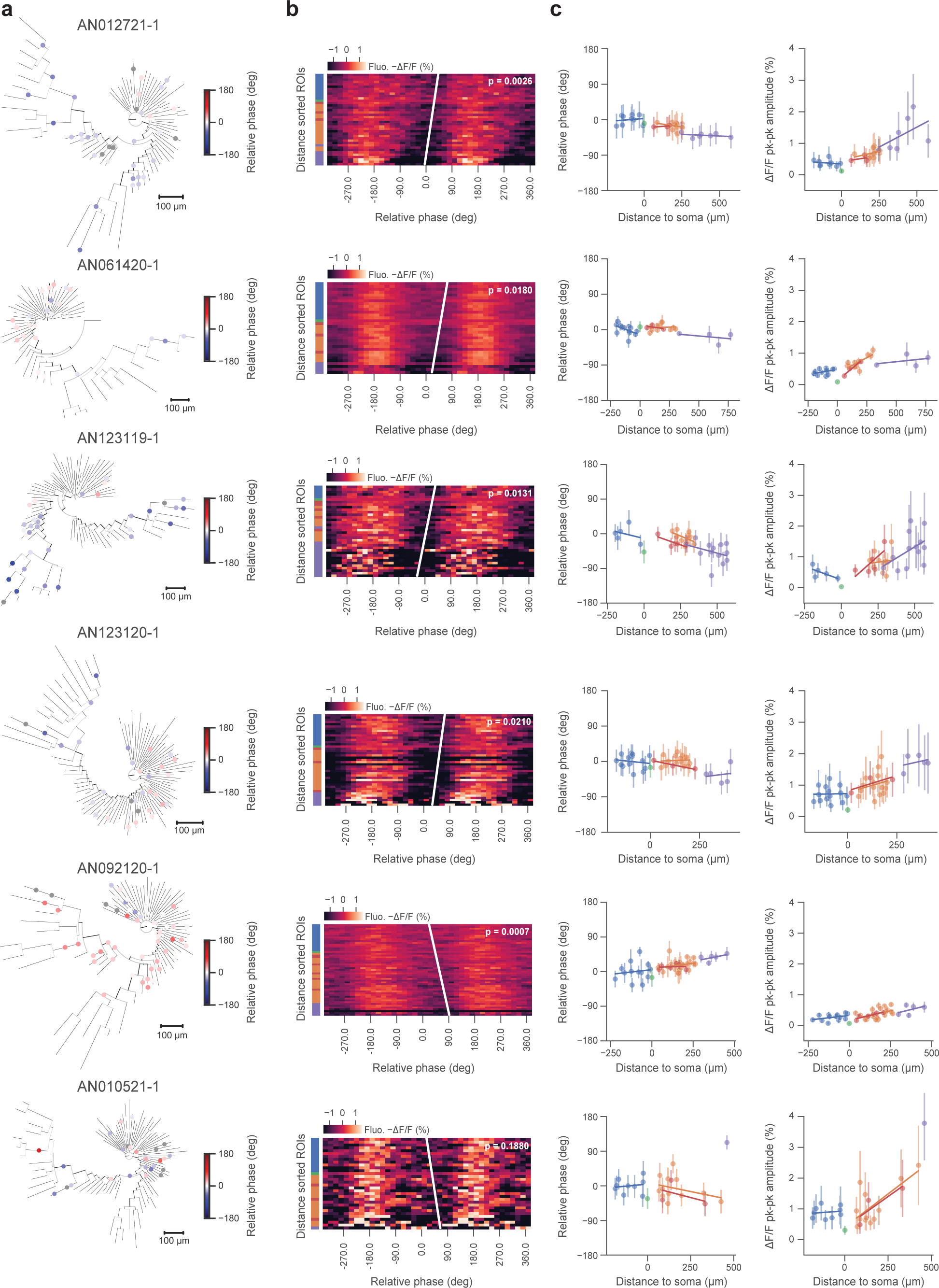
Extended data associated with Fig 2. Additional example cells. **a.** Morphological reconstructions with theta-oscillation modulated fluorescence ROIs colored by relative phase to somatic compartment. Nonsignificantly phase modulated ROIs in grey. **b.** Membrane potential oscillation gradients for cells shown in **a.**, with rank regression line and *p* value shown. **c.** Per-region phase relative to somatic compartment, amplitude estimates and associated trendline fits for cells shown in **a.**.

**Figure S9.**
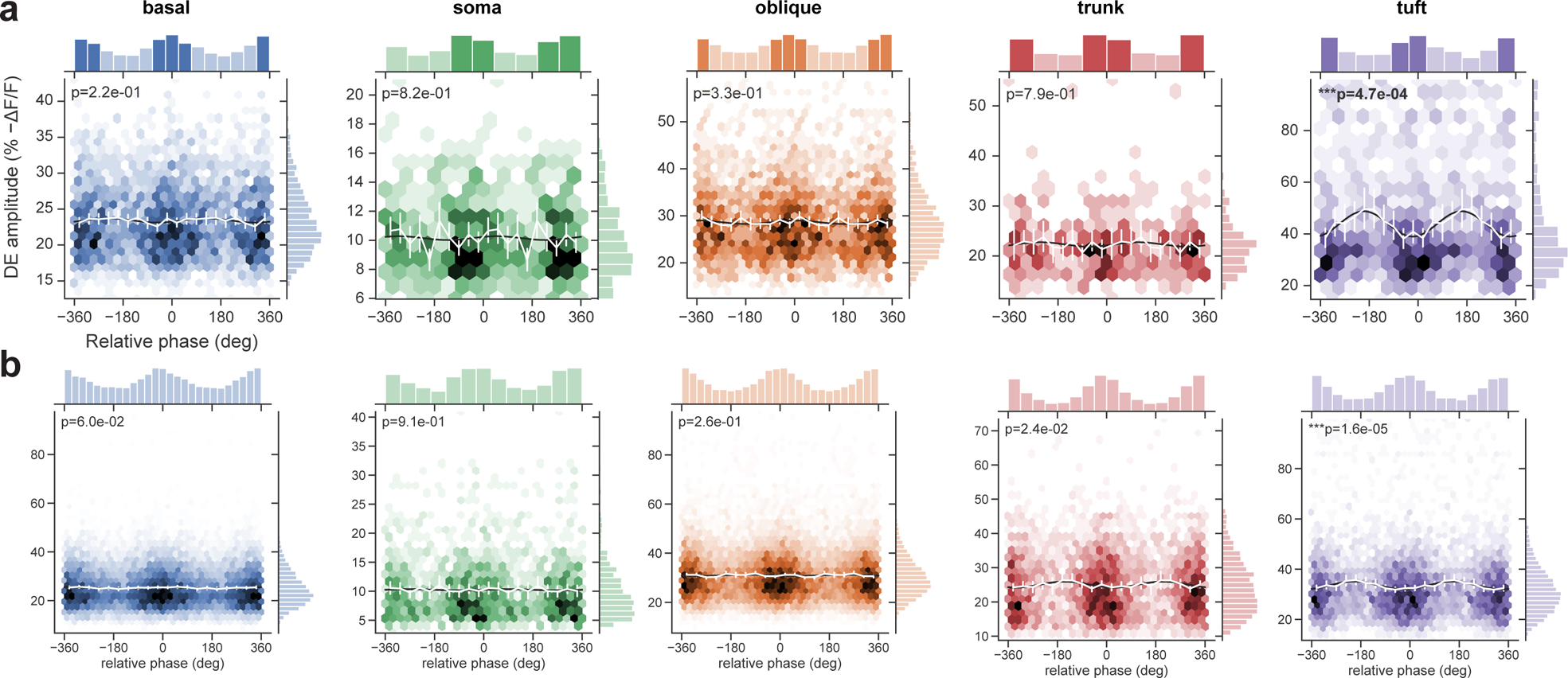
Extended data associated with Fig 2. Theta-phase modulation of DE amplitude across somatic and dendritic regions. 2D histograms of the distribution of DEs in theta phase and amplitude space. Each hexagon corresponds to a single phase/amplitude bin, while color intensity corresponds to the relative density of events within that bin. **a.** DEs from example cell AN092120-1 shown in Fig S8 and **b.** for all cells. Linear-circular fits (black lines; see Methods: Phase modulation of DE amplitude) are shown along with significance level and mean amplitude of DEs per phase bin (white lines) with 95% CI whiskers. Across cells, 2/11 cells showed DE amplitude modulation of at least one region (*α* = 0.01 using the Bonferroni correction).

**Figure S10.**
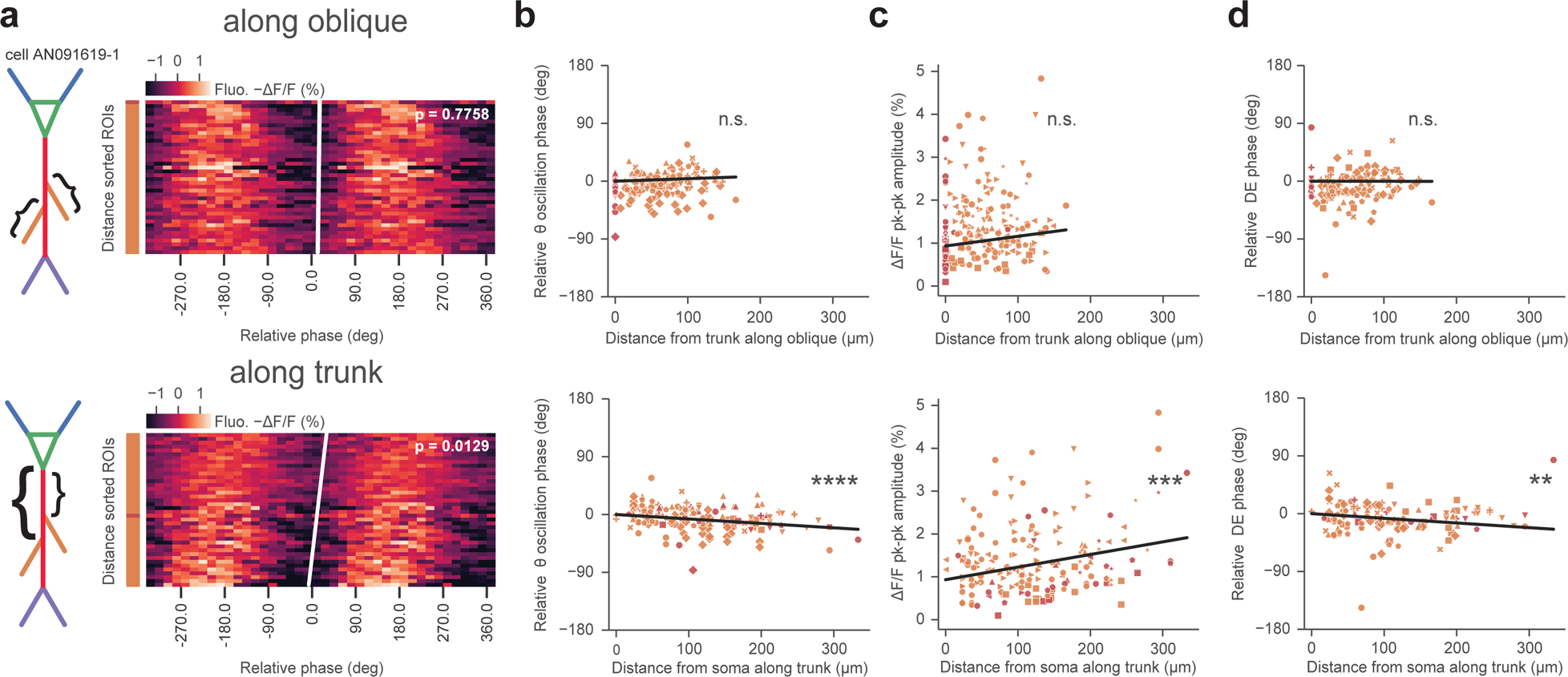
Extended data associated with Fig 2. Theta-band membrane potential oscillation phase, amplitude and DE phase gradients occur along trunk rather than along oblique dendrites. Phase measured relative to the actual or inferred somatic compartment phase. **a.** Theta-band phase-binned −Δ*F/F* fluorescence heatmaps of distance-sorted trunk and oblique dendritic ROIs for example cell of Figure 2 along with rank-regression lines, sorted by distance along oblique w.r.t. trunk stem location (top) or along trunk w.r.t. somatic compartment (bottom). Phase advances significantly (*p* = 0.0129) along trunk but not along oblique dendrites (*p* = 0.7758). Multivariate linear regression of theta-band relative fluorescence oscillation phase, −Δ*F/F* pk-pk oscillation amplitude and DE phase (**b.** oscillation phase: adjusted *r*^2^ = 0.18*, p <* 1.3 × 10^−7^, **c.** oscillation amplitude: adjusted *r*^2^ = 0.07*, p* = 0.002 and **d.** DE phase: adjusted *r*^2^ = 0.07*, p* = 0.002) has a nonsignificant projection against distance along oblique (top row, **b.** oscillation phase: *p* = 0.12, **c.** oscillation amplitude: *p* = 0.12, **d.** DE phase: *p* = 0.973) but significant along trunk (bottom row, **b.** oscillation phase: *p <* 1.2 × 10^−7^, **c.** oscillation amplitude: *p* = 0.0005, **d.** DE phase: *p* = 0.004) dendrites for ROIs pooled from all cells.

**Figure S11.**
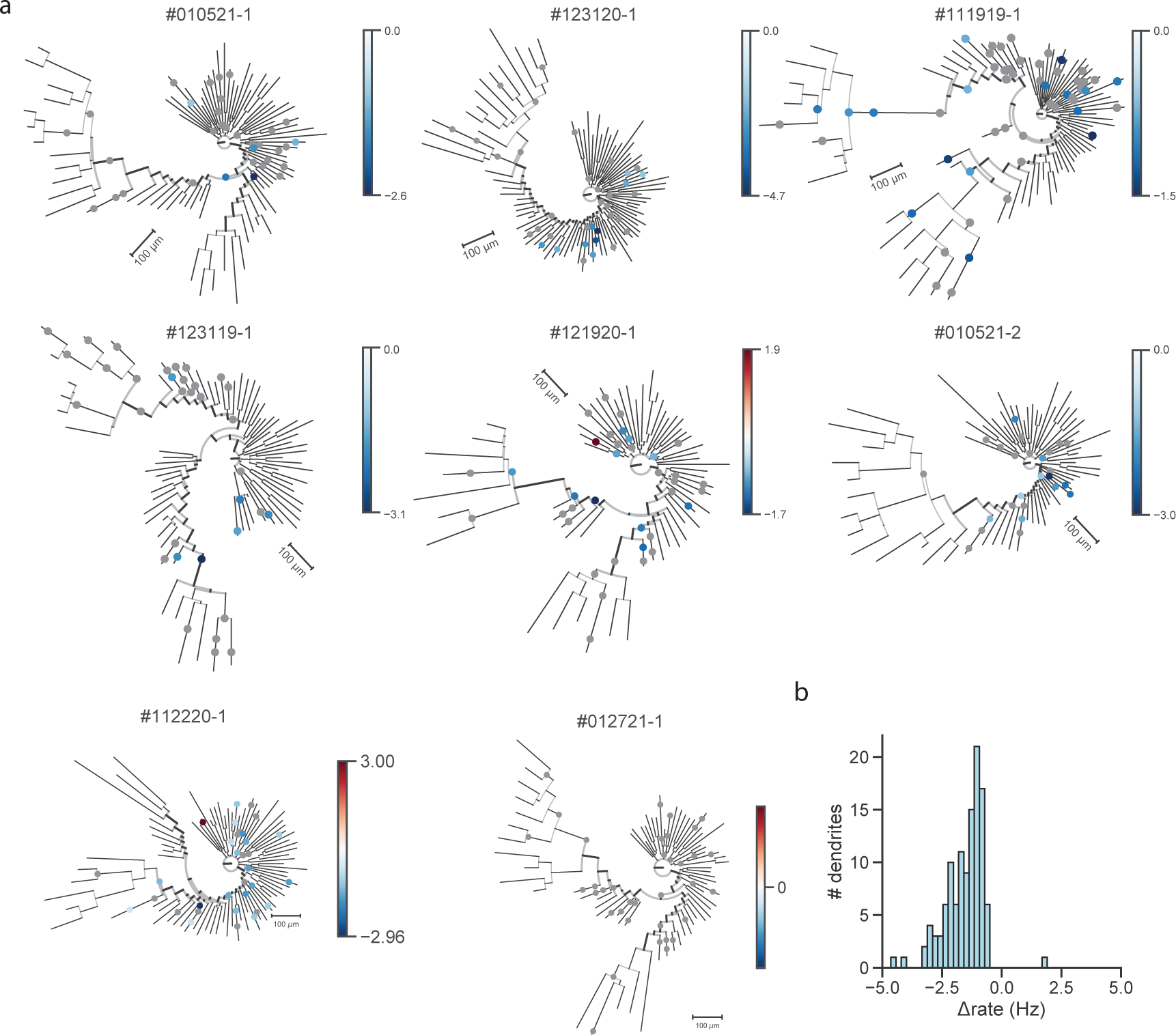
Rate modulation of individual dendrites by SWRs. **a.** Phylograms of all cells analyzed in Fig 4, with recording locations colored by change in DE rate inside minus outside of SWRs. Grey: Nonsignificantly ripple modulated locations under Poisson rate test. This test is generally conservative given the sparsity of events. 17.7% of dendrites exhibit significant rate modulation around ripples, with the majority suppressing their firing in response to SWRs. **b.** Summary histogram of rate modulation (*r_inside_ − r_outside_*) at significantly modulated dendrites

**Figure S12.**
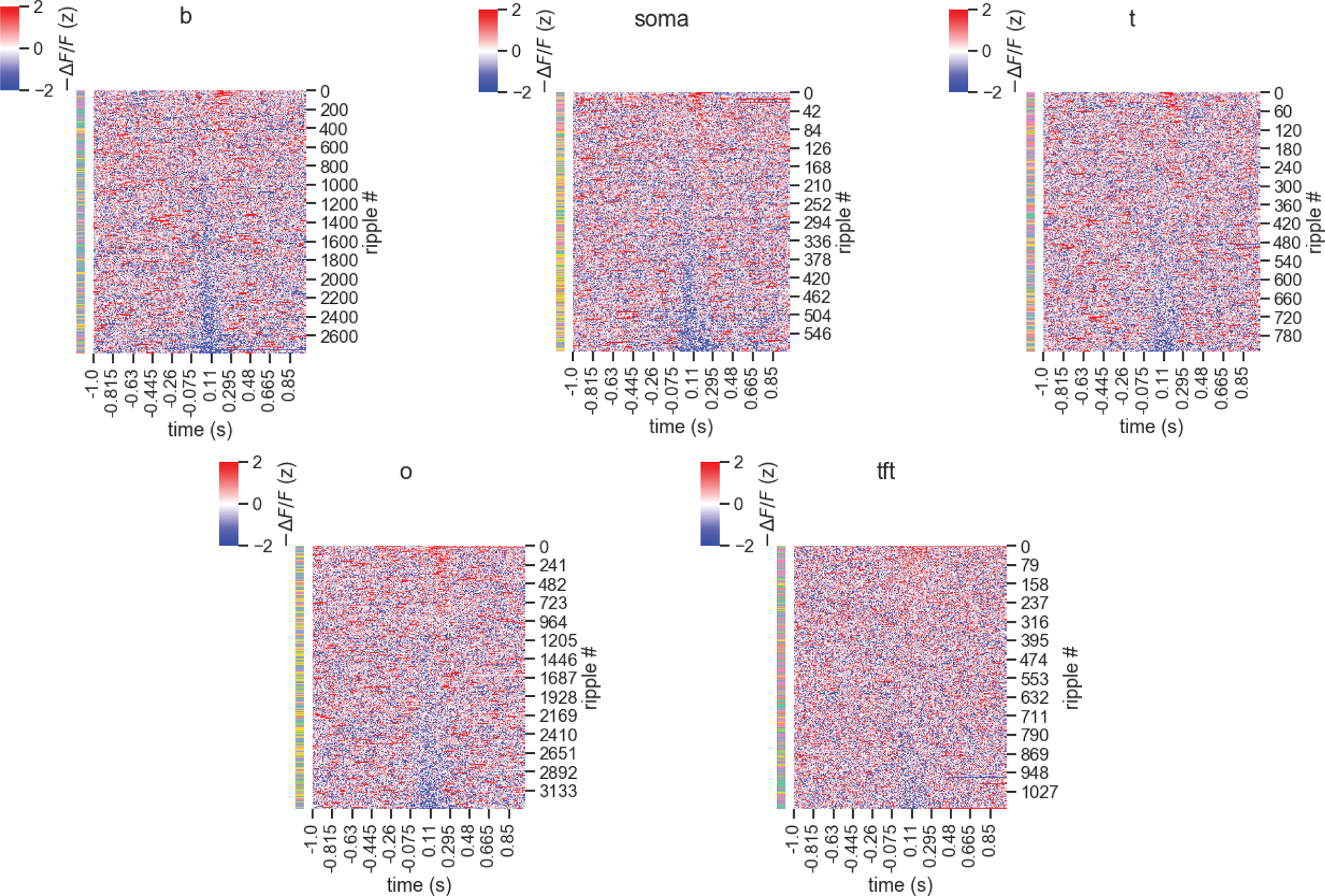
Heatmaps of all ripples shown in Fig 4. Heatmap of z-scored fluorescence around SWRs across all cells and segments by region, sorted by mean activity from 0-250 ms. Row color: cell ID. Inhibition predominates in the peri-ripple zone, but a small proportion of ripples recruit the recorded dendrite.

**Figure S13.**
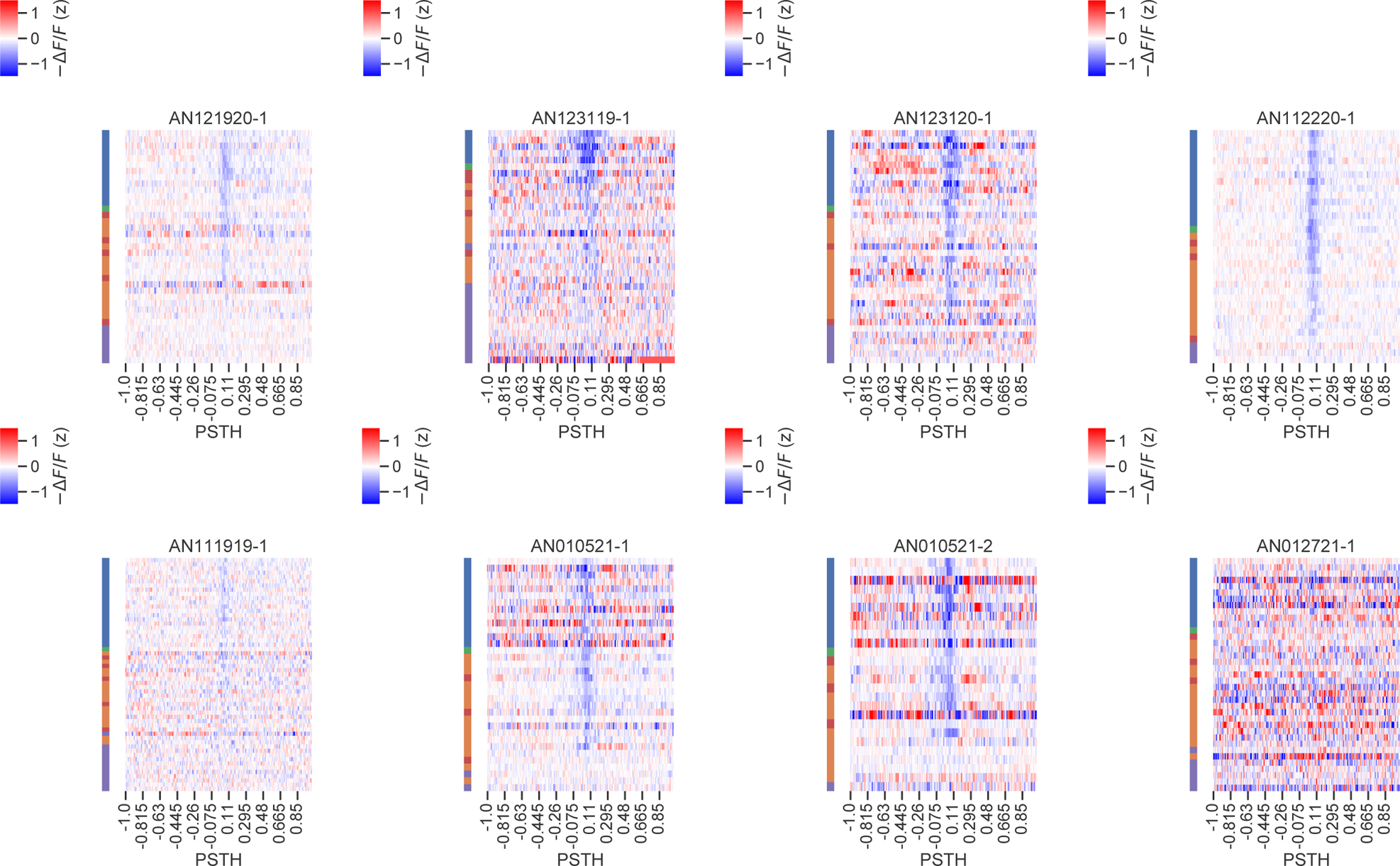
Dendritic hyperpolarization is centered around ripple peak irrespective of soma distance. Similar to our theta analysis of Figure 2, we ask the question of whether anatomical gradients emerge in the dendrite SWR response. Unlike our theta findings, it appears SWR-associated hyperpolarization is centered symmetrically about the ripple peak in all compartments. Mean heatmap of z-scored fluorescence in a ±1 s window around SWRs (aligned to peak) by cell and segment, sorted by signed soma distance. Row color: recorded region. Inhibition predominates in the peri-ripple zone in most cells and segments. Cell AN012721-1 did not exhibit strong ripple modulation.

**Figure S14.**
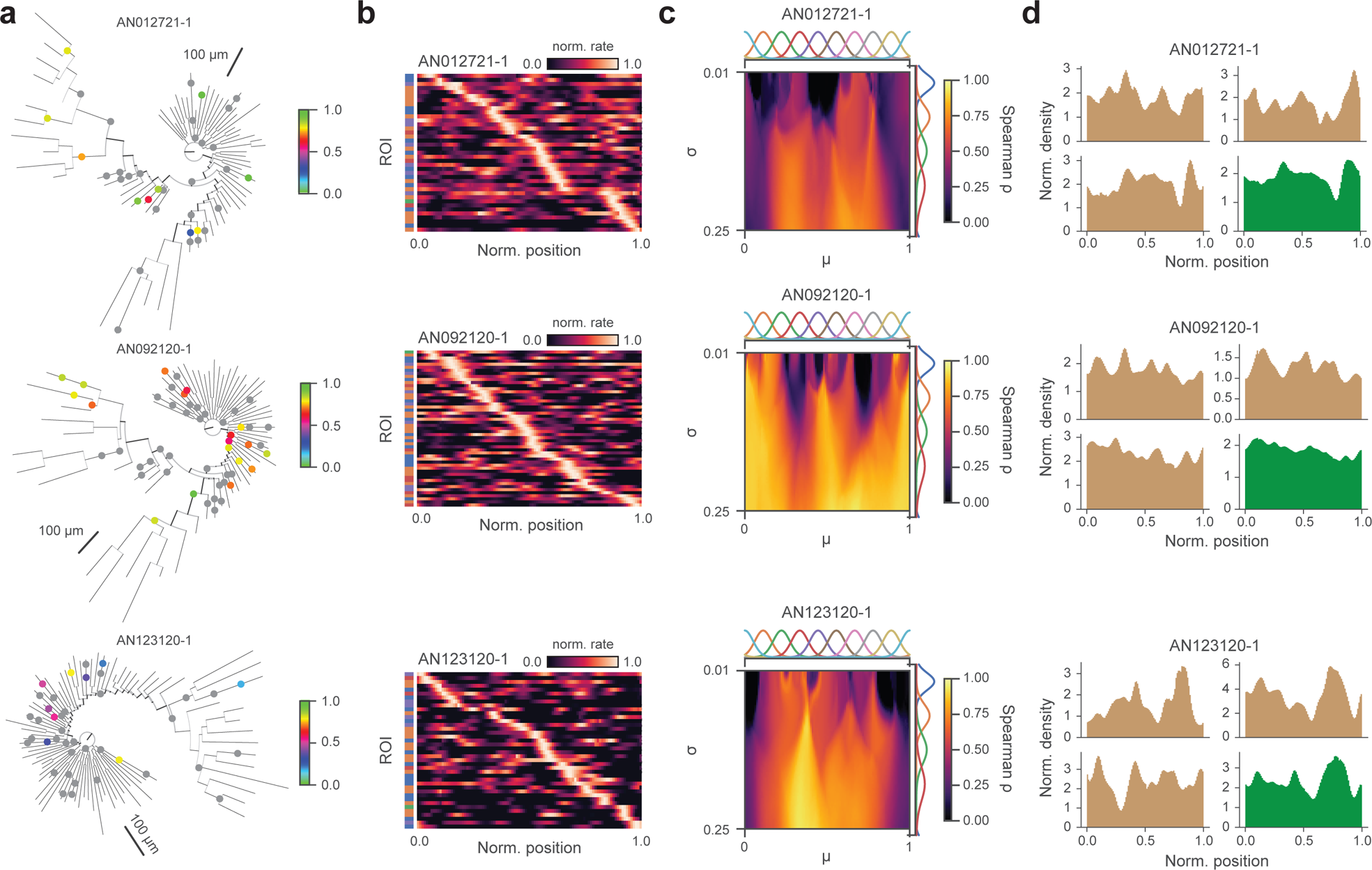
Extended data associated with Fig 5. We provide further examples of cells from our dataset exhibiting distributed dendritic spatial tuning. Similar to the example we show in the main figure, these cells feature diverse tuning across the dendritic arbor and high expressivity for constructing somatic tuning curves. **a.** Additional example cells, with recording locations colored by tuning centroid. Grey: Nonsignificantly peaked tuning curves under Rayleigh test. **b.** Segment tuning curves associated with cells in **a.**, sorted by tuning curve peak. **c.** Reconstruction fidelity matrix of Fig 5e for example cells shown. **d.** Tuning histograms of primary basal subtrees (brown), with overall basal tuning histogram (green).

**Table S1.**
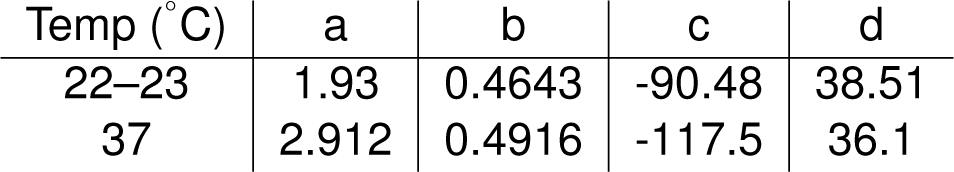
ASAP3 steady state fluorescence response in HEK293 cells as a function of membrane potential (in mV). Sigmoidal fit coefficients for *F* (*v*) = *a* + (*b − a*)/*{*1 + exp[(*c − v*)/*d*]}.

| Temp (°C) | a | b | c | d |
| --- | --- | --- | --- | --- |
| 22–23 | 1.93 | 0.4643 | -90.48 | 38.51 |
| 37 | 2.912 | 0.4916 | -117.5 | 36.1 |

**Table S2.**
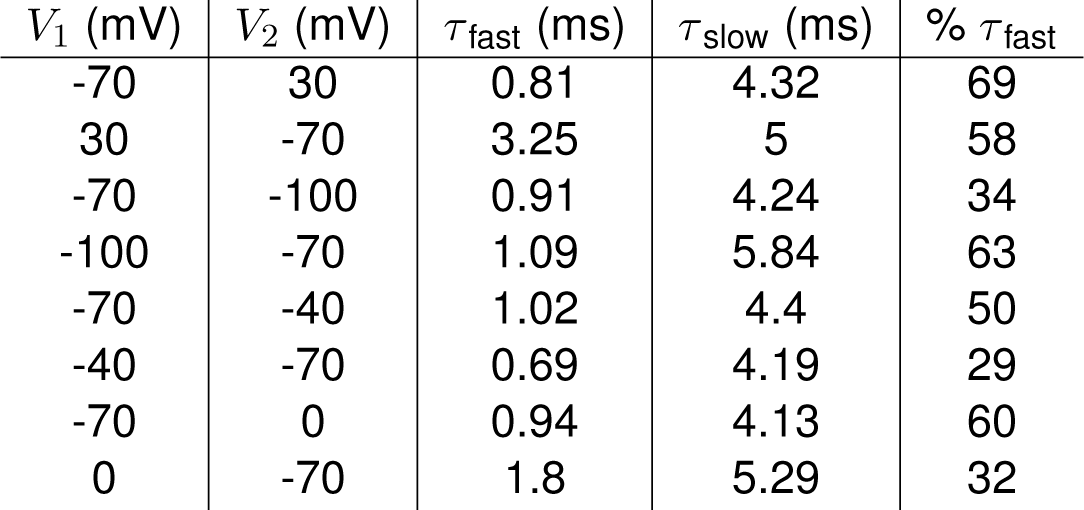
ASAP3 dynamic step double exponential fit response in HEK293 cells at 37 ^°^C for voltage steps *V*_1_ −*→ V*_2_.

| $V_1$ (mV) | $V_2$ (mV) | $\tau_{\text{fast}}$ (ms) | $\tau_{\text{slow}}$ (ms) | % $\tau_{\text{fast}}$ |
| --- | --- | --- | --- | --- |
| -70 | 30 | 0.81 | 4.32 | 69 |
| 30 | -70 | 3.25 | 5 | 58 |
| -70 | -100 | 0.91 | 4.24 | 34 |
| -100 | -70 | 1.09 | 5.84 | 63 |
| -70 | -40 | 1.02 | 4.4 | 50 |
| -40 | -70 | 0.69 | 4.19 | 29 |
| -70 | 0 | 0.94 | 4.13 | 60 |
| 0 | -70 | 1.8 | 5.29 | 32 |

**Table S3.** Data associated with Fig 1f. Median DE frequencies and 95% bootstrapped confidence intervals inside and outside running epochs (all values reported in Hz, *N* = 11 cells)

| region | run modulated | upper | lower | median |
| --- | --- | --- | --- | --- |
| basal | 0.0 | 1.40 | 1.06 | 1.25 |
|  | 1.0 | 0.74 | 0.63 | 0.68 |
| oblique | 0.0 | 1.66 | 1.34 | 1.55 |
|  | 1.0 | 0.87 | 0.71 | 0.82 |
| soma | 0.0 | 1.55 | 0.22 | 0.88 |
|  | 1.0 | 0.74 | 0.13 | 0.38 |
| trunk | 0.0 | 1.65 | 0.86 | 1.18 |
|  | 1.0 | 0.82 | 0.49 | 0.70 |
| tuft | 0.0 | 2.11 | 1.49 | 1.78 |
|  | 1.0 | 1.20 | 0.98 | 1.10 |

**Table S4.** Data associated with Fig 4d. Median DE peaks and 95% bootstrapped confidence intervals inside and outside SWR epochs (all values reported in % − Δ*F/F*)

| region | ripple | median | lower | upper |
| --- | --- | --- | --- | --- |
| basal | outside | 18.9 | 18.1 | 19.5 |
|  | inside | 15.7 | 13.9 | 18.3 |
| oblique | outside | 20.7 | 19.9 | 21.2 |
|  | inside | 18.9 | 16.8 | 21.6 |
| soma | outside | 6.7 | 3.9 | 8.6 |
|  | inside | 14.6 | n/a | n/a |
| trunk | outside | 16.7 | 15.4 | 17.5 |
|  | inside | 17.0 | 14.2 | 19.7 |
| tuft | outside | 18.8 | 16.5 | 19.5 |
|  | inside | 17.6 | 12.9 | 19.6 |

**Table S5.** Data associated with Fig 4e. Median delay and 95% bootstrapped confidence intervals between LFP SWR power peak vs membrane potential trough (all values reported in ms)

| region | median | lower | upper |
| --- | --- | --- | --- |
| basal | 40.60 | 0.0 | 57.21 |
| oblique | 65.27 | 5.91 | 77.44 |
| soma | 57.50 | 35.00 | 87.50 |
| trunk | 60.42 | 15.00 | 87.83 |
| tuft | 6.67 | -47.50 | 71.67 |

**Table S6.** Data associated with Fig 4f. Median DE frequencies and 95% bootstrapped confidence intervals inside and outside SWR epochs (all values reported in Hz)

| region | ripple | median | lower | upper |
| --- | --- | --- | --- | --- |
| basal | inside | 0.20 | 0.05 | 0.28 |
|  | outside | 1.04 | 0.89 | 1.61 |
| oblique | inside | 0.64 | 0.40 | 0.88 |
|  | outside | 1.38 | 1.14 | 1.68 |
| soma | inside | 0.0 | n/a | n/a |
|  | outside | 0.55 | 0.17 | 1.01 |
| trunk | inside | 0.30 | 0.0 | 0.58 |
|  | outside | 1.37 | 0.82 | 2.27 |
| tuft | inside | 0.72 | 0.24 | 1.50 |
|  | outside | 1.87 | 1.18 | 2.19 |

**Table S7.** Data associated with Fig 4h. Median membrane potential and 95% bootstrapped confidence intervals at LFP SWR power peak vs membrane potential trough (all values reported in % − Δ*F/F*)

| region | timepoint | median | lower | upper |
| --- | --- | --- | --- | --- |
| basal | LFP peak | -1.2 | -3.3 | -0.5 |
|  | fluor trough | -9.5 | -15.1 | -3.6 |
| oblique | LFP peak | -0.08 | -1.6 | 0.03 |
|  | fluor trough | -8.2 | -13.3 | -4.6 |
| soma | LFP peak | -0.4 | -0.8 | -0.2 |
|  | fluor trough | -1.4 | -2.0 | -0.7 |
| trunk | LFP peak | -0.4 | -1.3 | 0.1 |
|  | fluor trough | -4.8 | -7.3 | -2.5 |
| tuft | LFP peak | -0.5 | -3.6 | -0.1 |
|  | fluor trough | -12.4 | -16.3 | -6.2 |

**Table S8.** Data associated with Fig 4i. Median DE FWHM and 95% bootstrapped confidence intervals inside and outside SWR epochs (all values reported in ms.

| region | condition | upper | lower | median |
| --- | --- | --- | --- | --- |
| basal | nonripple | 31.92 | 20.73 | 25.00 |
|  | ripple | 42.04 | 10.17 | 27.31 |
| oblique | nonripple | 30.06 | 19.98 | 25.98 |
|  | ripple | 21.67 | 12.62 | 16.06 |
| trunk | nonripple | 33.25 | 16.38 | 28.00 |
|  | ripple | 35.18 | 3.04 | 17.03 |
| tuft | nonripple | 19.61 | 15.12 | 16.14 |
|  | ripple | 21.08 | 12.75 | 18.00 |

